# Wrapping it Up: Structural Basis of ADAMTS13 Global Latency

**DOI:** 10.1101/2025.11.26.689657

**Authors:** Norman Geist, Quintijn Bonnez, Karen Vanhoorelbeke, Mihaela Delcea

## Abstract

ADAMTS13 is a critical enzyme responsible for cleaving ultra-large von Willebrand factor (VWF) multimers, thereby preventing the formation of microthrombi in the microvasculature. Dysfunction or deficiency of ADAMTS13 is associated with thrombotic thrombocytopenic purpura (TTP), a rare and life-threatening disorder characterized by microangiopathic hemolytic anemia and severe thrombocytopenia. Understanding the regulation of ADAMTS13 is essential for developing therapeutic interventions for TTP. ADAMTS13 exhibits two latency mechanisms: local latency within the metalloprotease (MP) domain and global latency that affects its overall conformation. Although the role of the two CUB domains in binding the Spacer module is known during global latency, the contributions of TSP7, TSP8, and the flexible Linker region (L3) between TSP8 and CUB1 have been repeatedly observed but never explained. Binding studies with monoclonal antibodies (mAbs) targeting the MP domain have revealed a cryptic epitope that becomes accessible only when ADAMTS13 is conformationally activated, and Spacer-CUB binding is disrupted. However, the mechanism behind this long-range effect has never been explained. We present a novel autoinhibition model of ADAMTS13, proposing that its distal domains directly occlude substrate binding sites, thereby modulating the enzyme’s activity. In this model, TSP7 and TSP8 bind directly to the MP module, while the Linker region between TSP8 and CUB1 acts as a pseudosubstrate, mimicking the natural VWF-A2 substrate and blocking the binding sites. Our results show that the CUB domains must depart from the tandem arrangement observed in the crystal structure and instead adopt refined binding positions. Further evidence comes from novel monoclonal antibodies targeting more cryptic epitopes on various ADAMTS13 domains that we have recently presented, as well as pH- and EDTA-dependent conformational changes observed by ELISA. A detailed molecular model is derived from extensive molecular simulations and mechanistically reconciles a wide range of previously disconnected experimental observations. This novel insight into the regulation of ADAMTS13 lays the groundwork for new therapeutic strategies against TTP.

**Graphical Abstract:** 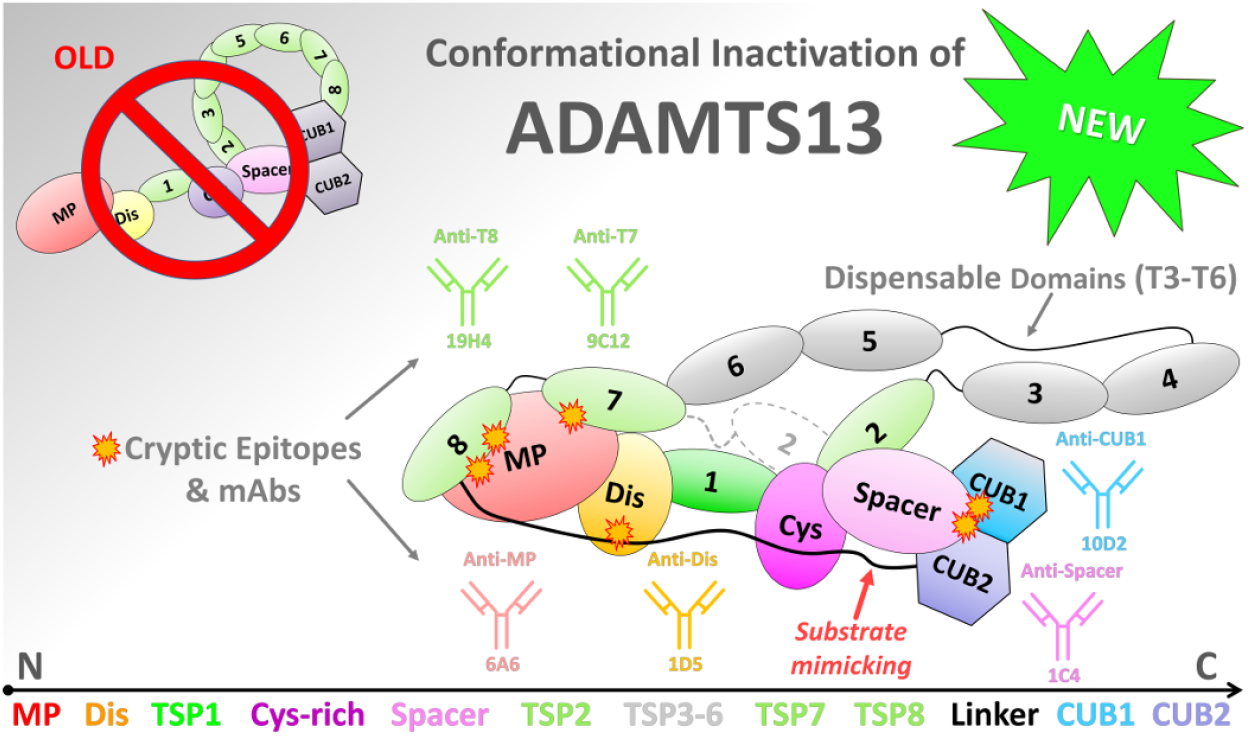

**Highlights:**

- Atomistic Structure of autoinhibited ADAMTS13-Del3To6
- L3 linker (TSP8–CUB1) has a dual function: acting as a pseudosubstrate that blocks VWF-A2 binding sites and guiding CUB1–2 positioning
- TSP7 and TSP8 bind at the MP domain to restrain activity
- CUB1 and CUB2 domains are not a single functional block

## Introduction

Thrombotic thrombocytopenic purpura (TTP) is a rare, life-threatening thrombotic microangiopathy caused by a severe deficiency of ADAMTS13 (A Disintegrin and Metalloprotease with Thrombospondin type 1 repeats-13) [1]. This deficiency can arise from biallelic mutations in the ADAMTS13 gene, leading to hereditary TTP (Upshaw–Schulman syndrome) [2, 3, 4], or from the development of anti-ADAMTS13 autoantibodies, resulting in immune-mediated TTP [1]. In both cases, insufficient ADAMTS13 activity allows ultra-large von Willebrand factor (VWF) multimers to accumulate, promoting microvascular thrombi and microangiopathic hemolytic anemia with severe thrombocytopenia [1].

Current treatments for hereditary TTP include plasma infusion and plasma exchange for immune-mediated TTP, immunosuppressive therapy and caplacizumap, reducing mortality from about 90% to 10%-20% [5, 6]. Recombinant ADAMTS13 has been proposed as an alternative therapeutic strategy [2, 7]. Understanding the molecular VWF recognition and interaction with ADAMTS13, as well as the enzyme’s inherent latency mechanisms, is crucial [8, 9, 10]. A comprehensive model of ADAMTS13’s auto-inhibition could serve as a structural platform for designing novel therapeutics not only for TTP but also for other diseases associated with disruptions in the ADAMTS13/VWF axis, such as myocardial infarction, stroke, and malaria[11, 12, 13].

ADAMTS13 regulates the size of VWF through proteolysis. Under shear stress, the A2 domain of VWF unfolds, exposing a specific binding and cleavage site for ADAMTS13. Several VWF-binding exosites, interacting with specific domains of ADAMTS13, are known to determine substrate specificity and play a role in two distinct allosteric mechanisms that activate ADAMTS13 in response to VWF (the “molecular zipper” model)[15]. ADAMTS13 is regulated via two independent forms of latency: a local latency state, in which key gatekeeper residues block substrate access to the metalloprotease (MP) active site, and a global latency state, maintained by binding of the distal CUB1–2 domains to the Spacer domain. Local latency is overcome due to the specific interactions with the natural substrate and co-factor: the unraveled VWF-A2 domain, also without any distal domains. The global latency state can be disrupted physiologically by binding of VWF D4–CK to ADAMTS13, displacing the CUBs from Spacer, or experimentally by anti-Spacer or anti-CUB monoclonal antibodies that disrupt the CUB–Spacer interaction, inducing conformational activation [8, 9, 10, 16]. This disruption leads to a 5-fold increase in ADAMTS13 catalytic activity [8, 16, 17, 18]. Interestingly, this increase in activity is not due to the exposure of the Spacer or Cys-rich domain exosites for VWF-A2. A substrate lacking the reciprocal binding regions for the Spacer or Cys-rich domains still leads to a similar activity increase when Spacer-CUB binding is disrupted. This suggests that Spacer-CUB binding affects the activity of the MP domain remotely, though the underlying mechanism for this allosteric effect is unresolved [8, 10, 19].

Modulation of ADAMTS13 activity requires not only the two C-terminal CUB domains, but also the TSP7, TSP8 repeats and the flexible Linker region (L3) between TSP8 and the CUB1-2 domains [16, 20]. In the following, we refer to the L3 region as “Linker”. The minimal ADAMTS13 construct retaining this full auto-inhibition can have TSP3 through TSP6 deleted, resembling the ADAMTS13 form found in pigeons [21, 22]. Under physiological conditions, global activation of ADAMTS13 is mediated by VWF-D4CK, which binds most strongly to the CUB1 domain but also interacts with TSP8 [10, 16], indicating an essential, yet unexplored role of TSP8 in the “global latency” mechanism. Binding studies employing anti-MP domain mAbs 6A6 and 3H9 have revealed a cryptic epitope in the MP domain [20, 23]. While 3H9 demonstrates equal binding affinity to both full-length ADAMTS13 and a truncated version of the MDTCS region (lacking distal domains), 6A6 exhibited a pronounced preference for binding MDTCS over full-length ADAMTS13. The same was observed in conformationally activated full-length ADAMTS13 [10]. Recent results from Hydrogen-Deuterium Exchange Mass Spectrometry (HDX-MS) revealed several surface regions that experience changes in their solvent accessibility during conformational activation, hinting at a larger conformational reassembly than previously anticipated [19, 24].

This study builds on new findings from mAbs targeting additional hidden epitopes on different ADAMTS13 domains[25] and integrates key experimental data to develop a unified conformational model of ADAMTS13’s auto-inhibition mode in circulation. We propose here that TSP7 and TSP8 bind directly to the MP domain, thereby restricting substrate access and limiting conformational freedom of ADAMTS13 (Figure 1, A). Additionally, a surprising sequence similarity is revealed between the Linker region and the VWF-A2 substrate. We identify the Linker region as a pseudosubstrate that bridges from TSP8 at the MP domain, across the VWF-A2 substrate-binding sites at the MP, Dis, and Cys-rich domains to the distant CUB1-2 domains, which then bind at Spacer.

**Figure 1:**
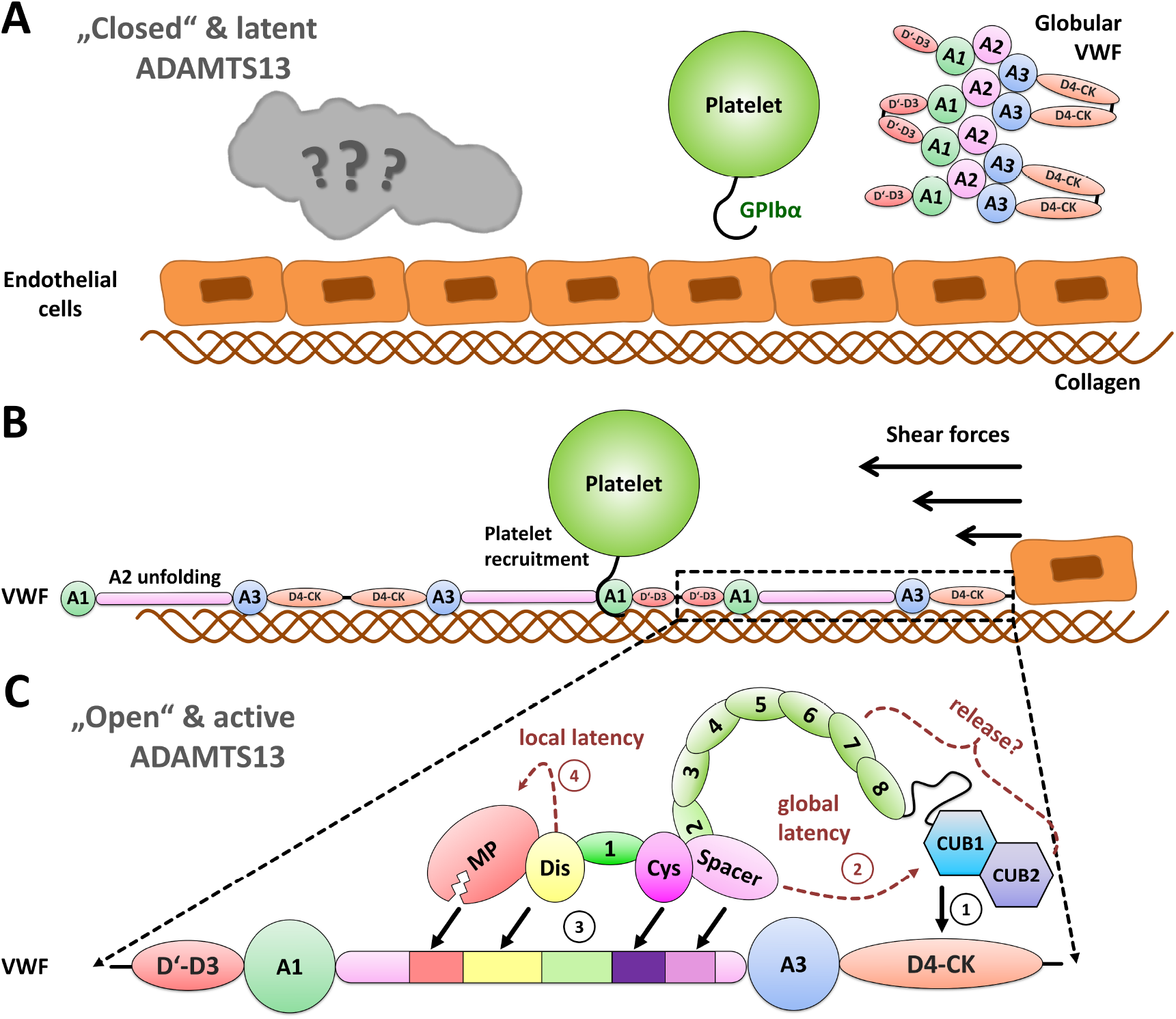
Currently proposed interplay of VWF and ADAMTS13. **(A)** In circulation, ADAMTS13 adopts a closed and inactive conformation (global latency) mediated by CUB1-2 binding to Spacer, with the exact mechanism unknown. VWF multimers are self-folded to prevent premature platelet binding. **(B)** Upon blood vessel damage, VWF-A3 binds subendothelial collagen, and shear stress unpacks the VWF multimers into extended chains, exposing the A1 domain that recruits platelets. The A2 domain completely unfolds into a linear peptide strand, exposing the cryptic cleavage site for ADAMTS13. **(C)** VWF D4-CK binding releases the CUB domains from Spacer **(1)**, also releasing activity restraints on the remote MP domain (global latency)**(2)**. The Spacer, Cys-rich, Dis and MP domains bind their exosites on the unraveled A2 domain, first in a precursor state **(3)**. The A2-strand acts as a cofactor and removes the gatekeeper tetrad[14] via an allosteric change that opens the active center (local latency) **(4)**. The MP domain can ultimately lock to the A2-cleavage site and cut the peptide strand.

Molecular dynamics simulations have recently provided structural insights into the VWF-A2 substrate binding[14]. There, we used our enhanced sampling TIGER2h_*P E*_[26] method to explore the large conformational space of the isolated ADAMTS13-MDTCS domains and their interaction with the unfolded VWF-A2 domain. In the present work, we used a similar approach and applied our TIGER2h_*P E*_ method to investigate the binding of TSP7 and TSP8 to the MP domain, the substrate-like interaction of the Linker region, and how the distal domains wrap around the MDTCS surface when the CUB1-2 domains are anchored to Spacer. This includes new binding modes for both CUB domains, which must dissociate from their recently observed fused structure during crystallization[9], leading to a cohesive structural “global latency” model, reconciling all available cornerstones from experimental groundwork. In order to minimize structural uncertainty, we applied a shortened ADAMTS13-Del3To6 construct, built with high confidence using AlphaFold2 (AF2) [27] based on available crystal structures. We further tested the dynamic nature of ADAMTS13 regulation by assessing its sensitivity to environmental and chemical factors such as pH and ion chelation.

## Results and Discussion

An initial structural model of ADAMTS13-Del3To6[21, 22], was generated using AF2 (Figure S1). The structure is conformationally active and the C-terminal CUB domains are not attached to Spacer. The Linker region appears linear and suspiciously resembles the unfolded VWF-A2 domain. We developed the final closed conformation of the ADAMTS13 construct through a series of refinement steps (Table S1).

### Binding sampling of TSP7 and TSP8 under natural constraints

We first conducted extensive structural sampling of a construct truncated after TSP8 (Sim1) to assess the conformations of TSP7-TSP8. Because these domains remain tethered to the MDTCS core through TSP2 at Spacer, their motion is limited to physiologically plausible binding poses (Figure S2). The simulations revealed several configurations in which TSP8 contacted MDTCS domains, supporting a preferred binding mode at the MP domain that would enable continuous connection to the Linker acting as a pseudo-substrate. Two such MP-associated poses were particularly compatible with positioning the Linker within the active-site cleft. In the larger cluster configuration (cluster2), the TSP8 module is attached to a helical region in the MP domain (L152-T160) by means of predominantly hydrophobic interactions (Figure S3). The interface at the TSP8 domain involved residues around I1092 and V1120-G1124. The TSP8 region W1126-C1130 interacted with both glycan chains at the MP domain. Further evaluation is required to ascertain whether this binding mode will also be found in the non-truncated model and the final closed form. Cluster4 showed a similar structure, with the interface on the MP domain exhibiting only a slight offset.

### Binding sampling of Linker pseudosubstrate

Literature suggests TSP8 must be in direct proximity to the MP domain to explain the cryptic epitope for mAb 6A6, which can only bind active ADAMTS13 or the MDTCS portion[20, 10, 25]. Besides the distance-spanning function, allowing the terminal CUB domains to bind at Spacer, while TSP8 would rest at MP, sequence comparison with ADAMTS13’s natural substrate VWF-A2 revealed a much more important role. At key binding positions, the sequence of the Linker region strongly resembles the amino acid composition of VWF-A2, yielding a total similarity of 31.37 % (Figure 2, A). Notably striking is the presence of distinct motifs, such as the **EE/DE** motif (E1142–E1143) near the Dis binding site, or the **PR** motif in the vicinity of the C-terminus (P1175-R1176). The MP binding sequence is almost identical but differs, particularly at the catalytic residue (P1141/M1605), which is naturally absent in the Linker, making it a textbook example for a pseudosubstrate, that will not induce the same reconfiguration of the gate-keeper tetrad[8, 14] necessary to activate the MP domain, concealing sequence variations that harbor the cofactor function of the VWF-A2 strand. A potential TSP1 binding region identified for VWF-A2 in our previous work[14] is completely missing on the Linker, hinting at a similar yet different binding mode. Beyond the Cys-rich binding site, the Linker shows no further similarity, indicating that it does not specifically engage Spacer, which is expected to be occupied by CUB1–2 during autoinhibition. This contrasts with VWF-A2, which contains a dedicated Spacer-binding sequence and interacts only after ADAMTS13 is activated and the CUB domains have detached.

**Figure 2:**
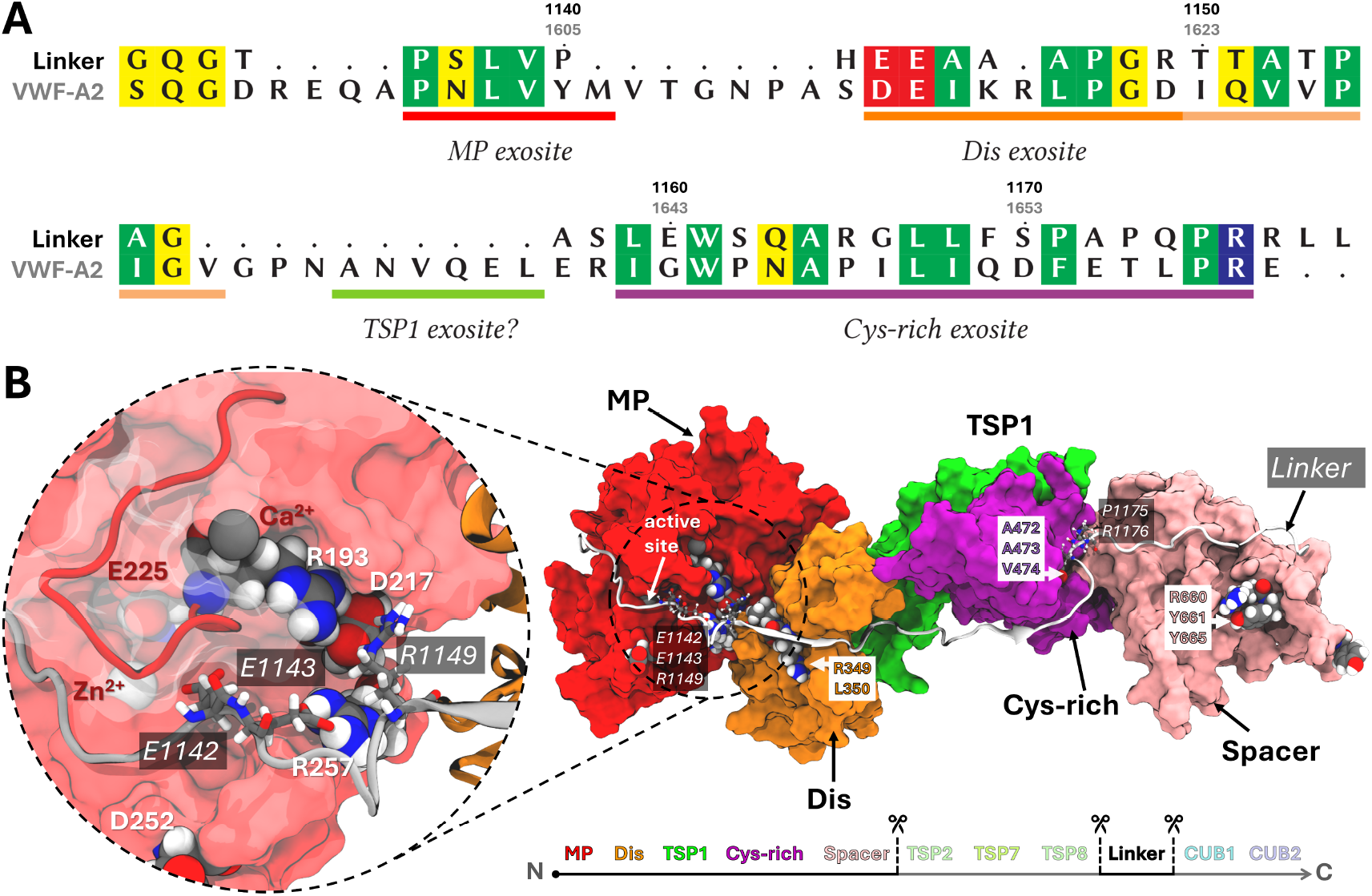
Sequence similarity between Linker pseudosubstrate and natural substrate for ADAMTS13: VWF-A2. **(A)** Sequence alignment between VWF-A2 and the Linker region. The previously identified binding sites of the A2 domain for interaction with specific ADAMTS13 domains are highlighted and show remarkable sequence similarity at these crucial positions. Correlating positions are colored according to their chemical properties: acidic negative (red), basic positive (blue), polar uncharged (yellow), and hydrophobic nonpolar (green). **(B)** Major cluster (0) resulting from Sim2 and RMSD cluster analysis shows binding in compliance with VWF-A2 at the Dis and Cys-rich modules of ADAMTS13. The prominent **EE** motif of the Linker (E1142-E1143) shows strong interactions with the gatekeeper tetrad of the MP domain. Another prominent motif **PR** (P1175-R1176), also present in VWF-A2, shows interactions at a proposed interaction site on the Cys-rich module.

We performed binding simulations for the Linker region against the MDTCS domains, using a similar approach to our study on VWF-A2 binding. Upon examining the structures, two noteworthy phenomena were identified: Initially, the **EE** motif on the Linker (E1142–E1143) appeared to interact and form a hydrogen-bond network with the gatekeeper tetrad residues of the MP domain, including several salt-bridges. The boundary between MP and Dis binding regions on the Linker is not as pronounced as for VWF-A2, given also several deletions between these regions (Figure 2, A). Secondly, the prominent **PR** motif (P1175–R1176) appeared to be involved in binding to the originally proposed Cys-rich exosite for VWF-A2 (A472–V473), a potential interaction not confirmed in our previous work. There, we identified the main passage interface of VWF-A2 on the rear side of the Cys-rich domain (I495-K497), while the originally proposed site (A472–V473) contacted VWF-A2 only rarely. In contrast, the Linker engages this site much more frequently, suggesting that the **PR** motif—previously linked to Spacer binding, may instead target the Cys-rich domain. This is likely reflected by the absence of a Spacer exosite for the Linker, allowing the motif to interact with its Cys-rich domain without competing restraints. A binding that may have been disfavored in our earlier VWF-A2 simulations, which relied on limited information about how specific A2 regions connect to ADAMTS13 domains.

Within the major structure identified (cluster0), significant interactions between the Linker and the gatekeeper tetrad include salt bridges E1142↔R193 and R1149↔D217 (Figure S4). The **EE** motif is further stabilized between R193, Q191, and V192; while E1143 remains anchored within the gatekeeper region, E1142 is also attracted toward the active-site zinc ion. Additional anchoring is contributed by the conserved region adjacent to the MP active site (Linker residues 1136–1139 and the corresponding 1601– 1604 segment in VWF-A2). Although the Linker occupies the same general pockets as VWF-A2[14], differences arise from the absence of the local helical motif, likely due to a sequence variation (S → N), and from the lack of enforced coordination between the analog carbonyl oxygen and the zinc ion. Consequently, the Linker motif adopts a slightly shifted position relative to the final VWF-A2 conformation.

There, the **DE** motif (D1614–E1615), and in particular the glutamic acid residue, was responsible for binding the Dis domain at position R349. However, in the Linker, several deletions between the MP- and Dis-binding regions result in the corresponding motif (E1142–E1143) shifted closer to the MP domain. In our present simulations, this motif integrated into a network with the gatekeeper tetrad. As a result, a different segment of the Linker engages the Dis domain, thereby forming a series of specific contacts (Figure S5). The binding pose is reminiscent of VWF-A2, including attachment to the *β*-sheet around S345– L351, with key interactions formed between R1149↔L351, T1151↔R349, T1153↔C347, and A1155↔S345. As observed for VWF-A2, the interaction continues on the rear side of the Dis domain around the *β*-sheet V320–F324, involving G1156↔F324, A1157↔T323, and additional diffuse contacts.

For the Cys-rich module, the binding also closely resembled that of VWF-A2, including *β*-sheet attachment around I495–K497 (Figure S6). Major contacts are built between L1167↔M496, L1168↔I495, F1169↔F494, S1170↔S493. This time also stable interactions are featured with the earlier proposed exosite around A472–V474, including several hydrophobic contacts, including: A1172↔H476, Q1174↔A472, P1175↔P475, and further contacts around L443, indicating an interface to harbor the **PR** motif (P1175–R1176).

Another notable effect of Linker binding to the ADAMTS13-MDTCS domains is an increase in direct contacts between the Dis and Cys-rich modules (Figure S7), compared to the VWF-A2. In the current Sim2, close contacts (<3.3 Å) were present in 38% of the generated states, whereas in Simulations 2–4 of our earlier work, they occurred in only 6%–7.5% of states. This difference is likely due to the ten–amino acid deletion between the Dis- and Cysrich–binding segments of the Linker, which may promote a more compact arrangement of ADAMTS13 in its latent state.

### Binding sampling of CUB1 and CUB2

We combined the structures of the TSP8 module bound to the MP domain (Sim1) with a Linker-binding pose that allowed seamless connection and extension by the terminal CUB1 and CUB2 domains (Sim2), allowing further sampling of their interactions at Spacer (Sim3). This unified model also allowed us to further monitor the behavior of TSP7, TSP8, and the Linker, which showed a strong tendency to pack against the MDTCS core, thereby reducing the accessible surface area. In particular, TSP7 attached to the MP–Dis interface and the two CUB domains explored multiple binding configurations on Spacer, although none matched the interface described by Kim *et al*.[9]

From Sim3, we selected a representative state with TSP7 bound to the MP–Dis interface to initiate further sampling (Sim4), during which TSP7 attached even more tightly to the MP and Dis modules. The Linker region adopted binding states strongly consistent with those observed in Sim2, again showing frequent engagement of the **PR** motif close to the originally proposed Cys-rich exosite (A472– V473). However, as in Sim3, numerous binding poses were sampled for the terminal CUB1-2 domains at Spacer, but none corresponded to the interface described before.

We therefore revised our approach and examined the CUB1 and CUB2 domains separately, rather than in their fused arrangement as seen in the crystal structure (PDB ID: 7B01). Using AF2, we predicted the binding of each CUB domain to a construct of Cys-rich and Spacer. For CUB2, no plausible binding states were obtained, consistent with the observations of South *et al*.[16], who showed that although both CUB domains are capable of binding an ADAMTS13-MDTCS construct, CUB1 binds more strongly and only CUB1 inhibits proteolysis. In contrast, AF2 produced robust and consistent predictions for CUB1. In three out of five models returned, the binding configuration was identical (Figure 3, A) and closely matched the knockout results reported by Kim *et al*.[9], involving the CUB1 interface residues W1245, W1250, K1252, and K1265. Singular mutations at these sites were sufficient to reduce CUB1 binding and abolish the effect of activating mAbs (Figure 3, B). The Spacer exosite residues R660, Y661, and Y665, proposed to interact with the VWF-A2 substrate, were found to engage directly with CUB1, providing a mechanistic explanation for its inhibitory effect on proteolysis. An overlay with the crystal structure of the fused CUB domains further showed that the observed CUB1 binding mode would require release of CUB2 from the tandem arrangement (Figure 3, C). These findings strongly support that we have identified the correct binding mode for CUB1, one that is inaccessible when CUB1 and CUB2 remain fused, and modestly distinct from the docking results reported by Kim *et al*.[9].

**Figure 3:**
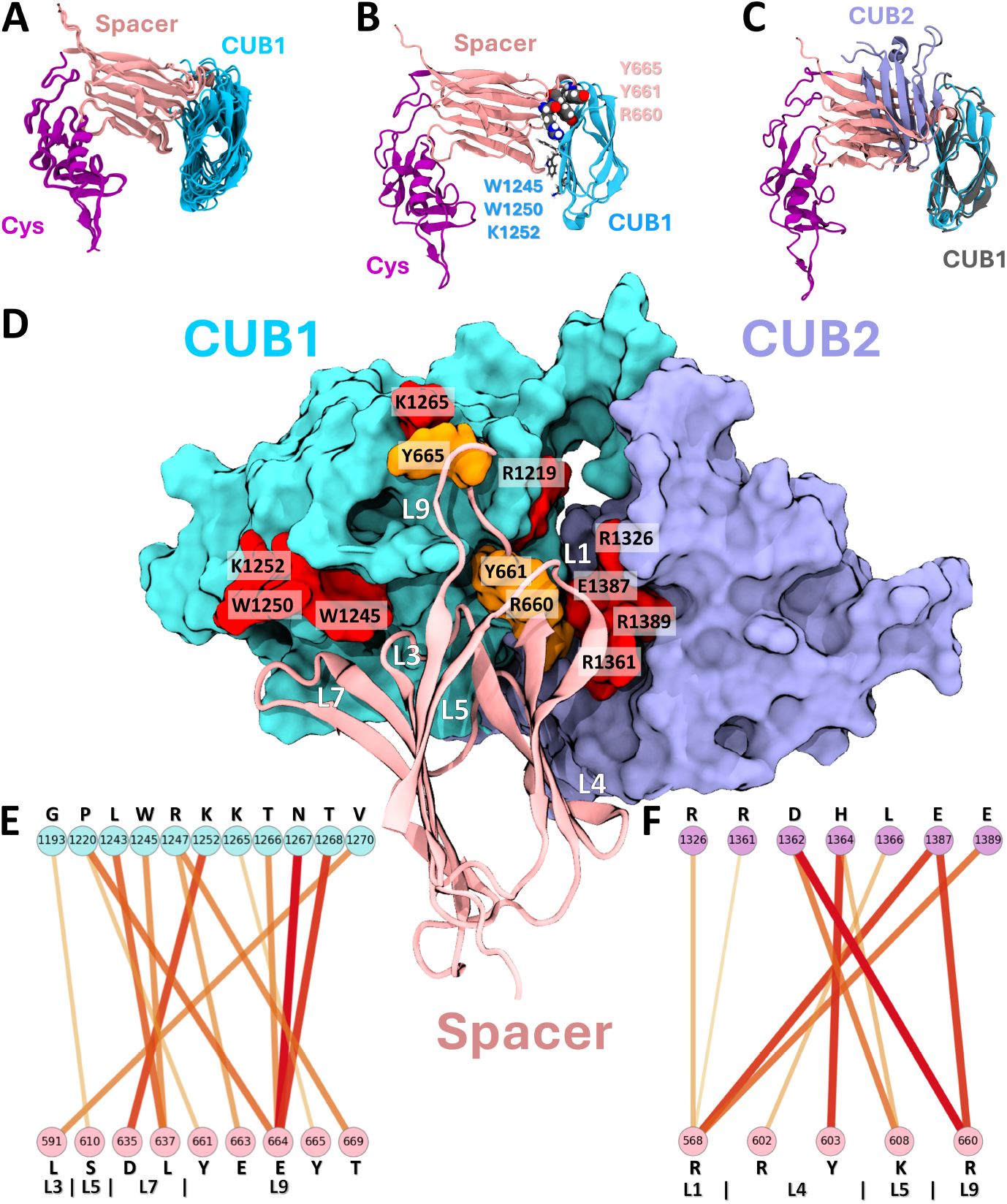
Novel binding positions of CUB1 and CUB2 on Spacer. **(A)** Structural overlay of three similar AlphaFold2 (AF2) models for CUB1 on Cys-rich and Spacer. **(B)** Best AF2 model with important residues highlighted. **(C)** Best AF2 prediction in overlay with crystal structure of CUB1-2 (PDB ID: 7B01) illustrating interference of CUB2 in this fused tertiary state. **(D)** Final binding positions of CUB1 and CUB2 at Spacer obtained through Sim5 (cluster0), with involved loops numbered (white). Red surface residues on CUB1-2 denote crucial positions found by Kim *et al*.[9], including R1219. **(E)** Residue interaction network of CUB1 domain (cyan) and Spacer (pink) with amino acid information and loop assignment. Contacts of affinities below 0.5 were cut off (full networks in SI). Connection line thickness and transparency quadratically weighted by affinities and colored from red to white for high and low affinity values. **(F)** Residue interaction network of CUB2 domain (plum) with Spacer (pink) as previously.

In the final simulation (Sim5), we hence combined a representative Sim4 state, containing the established TSP7, TSP8, and Linker interactions, with an additional restraint guiding CUB1 toward the AF2 predicted configuration. With most distal domains already positioned, the CUB domains were now free to explore their binding poses independently. This simulation produced well-defined binding modes for both CUB1 and CUB2 while simultaneously refining the inter-actions of the remaining distal domains with the MDTCS core. The resulting CUB1–2 arrangements in the major structural state (cluster0), together with the coherent overall packing, constitute the final autoinhibited conformation of ADAMTS13-Del3To6 (Figure 4, Figure S8, Figure S9).

**Figure 4:**
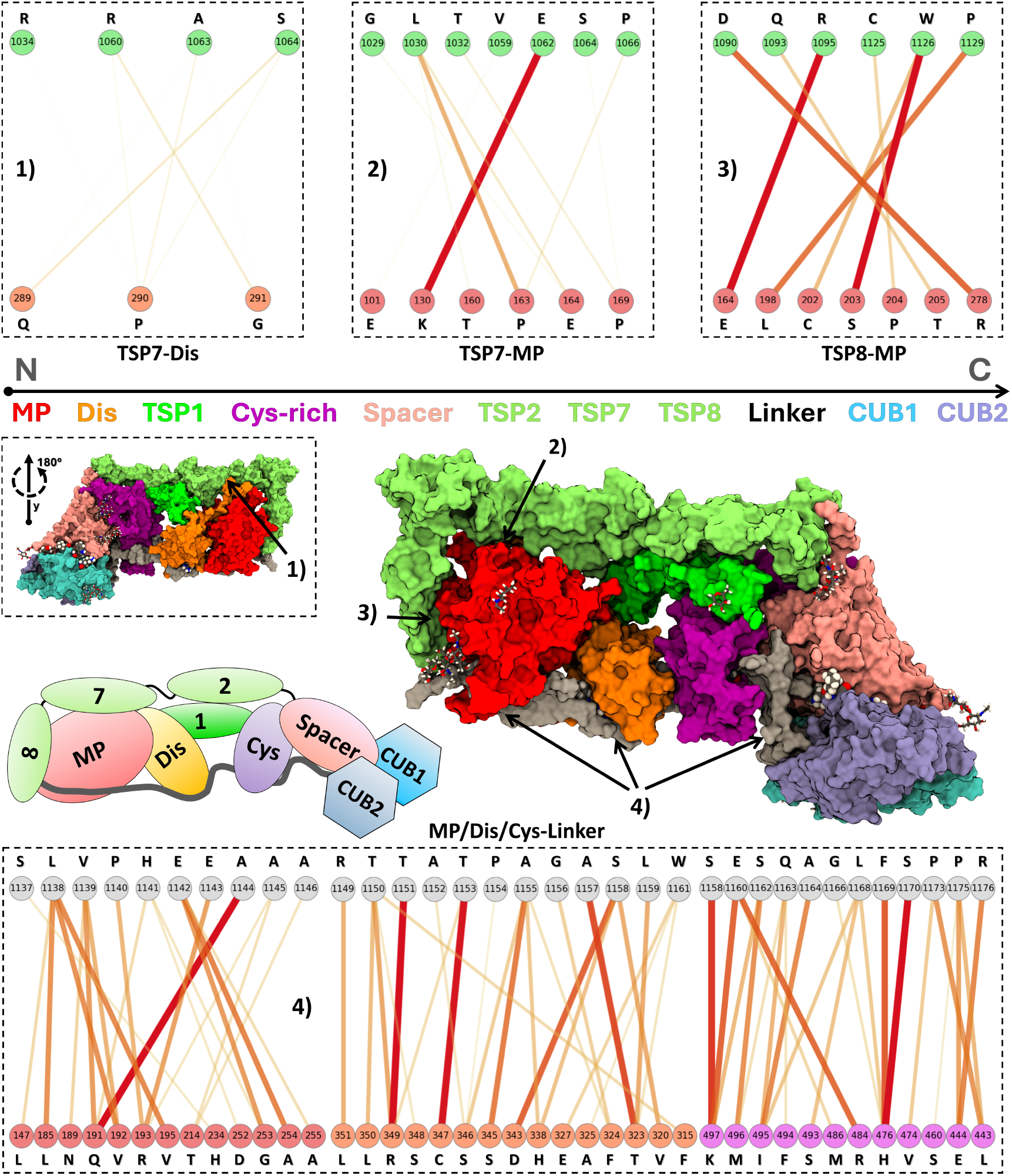
Final conformationally inactive ADAMTS13-Del3To6 model. Surface representation of ADAMTS13-Del3to6, illustrating compact wrapping of distal domains in the latent state. Sequential order and color scheme of domains are indicated. TSP7 and TSP8 are tightly associated with the MP domain and partially with the Dis domain. Linker spans and binds substrate-binding sites of MP, Dis, and Cys-rich modules. Both CUB1 and CUB2 engage Spacer. Inset shows structure rotated by 180^°^ around Y-axis. Residue interaction networks at different interfaces are annotated by numbers **(1-4)**. Connection line thickness and transparency quadratically weighted by affinities and colored from red to white for high and low affinity values. Contacts of affinities below 0.5 were cut off (full networks in SI). Schematic shows overall structure representation and domain organization.

### Autoinhibited conformation of ADAMTS13-Del3To6

The CUB1 domain largely maintained the arrangement predicted by AF2, engaging directly with the critical residues K1245, W1245, and W1250 (Figure 3, D–E, Figure S10). Together with nearby residues, these form a binding pocket for the L7 loop of the Spacer domain. Specific interactions included a salt bridge between D635↔K1252 and hydrophobic contacts involving L637↔L1243/W1245. The L3 loop contributes an additional contact between L591↔V1270. The L9 loop, containing residues R660, Y661, and Y665, participates in extensive interactions with CUB1, including a salt bridge between E663↔R1247 and further interactions with E664↔T1266/N1267/T1268, and Y665↔K1265. The importance of K1265 in CUB1 binding was previously highlighted by Kim *et al*.[9]. In addition, they reported R1219 as critical for disrupting CUB1 binding, though its role was unclear at the time because R1219 is positioned on the opposite side of the fused CUB1–2 structure. In our final configuration, however, R1219 clearly contributes to the Spacer-binding interface once CUB2 is free to move independently. This position has been reported previously to abolish proper secretion when mutated as R1219Q[28], while R1219W mutations are associated with abolished secretion in cTTP reports[29]. Notably, the two CUB domains cooperatively “sandwich” the L9 and L1 loops of Spacer. CUB2 engages strongly with the L9 loop, forming a salt bridge between R660↔D1362/E1387 (Figure 3, F, Figure S11). Additional contacts include a salt bridge between K608↔D1362 (L5), as well as hydrophobic interactions with L4, the strongest being Y603↔H1364. For the L1 loop, CUB2 establishes a strong salt bridge between R568↔E1387/E1398, interactions consistent with residues previously identified as critical for CUB2 binding.

The distal domains wrapped compactly around the MDTCS core, with TSP7 establishing only weak contacts with the disordered C-terminal tail of the Dis domain, running across the backside of MP (Figure 4 1, Figure S12), but forming stronger and more stable interactions directly with MP. These included a salt bridge between K130↔E1062 and hydrophobic contacts P163↔L1030/P1066 and P169↔T1032 (Figure 4 2, Figure S13).

TSP8 displayed a more extensive interface with MP compared to TSP7. While its binding overlapped with the interfaces observed in Sim1, the domain slightly shifted and rotated, becoming more firmly locked in place due to simultaneous engagement of the Linker and TSP7. Key stabilizing contacts included salt bridges E164↔R1095 and R278↔D1090, a hydrogen bond from S203 to the backbone of W1126, and hydrophobic interactions L198↔P1129, C202↔W1126, P204↔C1125, as well as T205↔Q1093 (Figure 4 3, Figure S14).

The Linker between TSP8 and CUB1 adopted a binding mode broadly consistent with Sim2 (MDTCS + Linker only), continuing to occupy the MP active center and blocking substrate access (Figure 4 4). However, some differences in detail were noted. In Sim2, guiding restraints were applied to reproduce VWF-A2 binding within the active site cleft, whereas in Sim5 the presence of TSP8 may have altered the final binding configuration. Specifically, although a salt bridge from E1143 of the **EE** motif to R193 of the gatekeeper tetrad was preserved (Figure S15), the remaining gatekeeper residues no longer directly engaged the Linker. Instead, R1149 and the E1142–E1143 pair stabilized a hairpin structure between MP and Dis, which appeared to obstruct access to the catalytic site. These deviations may represent one of multiple Linker binding modes observed in Sim5, with the extracted CUB1–2 cluster capturing only one preferred configuration. It is therefore proposed that both binding modes should be considered physiologically relevant, potentially corresponding to different stages during ADAMTS13’s autoinhibition. Several patient studies have reported that substitutions of R193 by tryptophan or glutamine causes a dramatic reduction in ADAMTS13 secretion and activity[30, 31]. While this has often been attributed to misfolding, our structural data suggest that these mutations may also disrupt a crucial electrostatic node required for proper domain packing and maintenance of global latency. Loss of R193-mediated stabilization could destabilize the MP–Linker interface, preventing correct autoinhibition during biosynthesis and leading to ER retention or degradation. Similarly, mutations such as R1219W in the CUB1–Spacer interface may act through a comparable mechanism, compromising the autoinhibited conformation of ADAMTS13 and thus failing quality control during secretion. Furthermore, stronger involvements of Linker residues A1144 and A1146 with the MP domain were observed. These positons were previously reported to disrupt ADAMTS13’s autoinhibition when mutated to Valine[32] and are currently the only known mutational experiments done to the Linker.

In contrast, Linker interactions with the Dis domain were essentially unchanged compared to Sim2, remaining tightly locked onto two *β*-sheets, consistent with our previous observations for both Sim2 and VWF-A2 binding (Figure S16).

At the Cys-rich module, the binding pose shifted slightly relative to Sim2, altering some residue-level interactions. Nonetheless, the **PR** motif (P1175–R1176) remained engaged near the previously proposed exosite around A472–V474, primarily through hydrophobic contacts (Figure S17). These variations may again reflect either alternative binding modes by chance coexisting with the CUB1–2 domain arrangements identified in clustering, or conformational changes introduced by the presence of the CUB domains and the overall compact packing of Dis, Cys-rich, Spacer, and CUB1–2.

### Conformational autoinhibition mechanism

Further support for our model comes from our identification of additional mAbs targeting cryptic epitopes (6A6 against MP domain, 1D5 against Dis domain, 1C4 against Spacer domain, 9C12 against TSP7 domain, 19H4 against TSP8 domain, and 10D2 against CUB1 domain) in circulating ADAMTS13, all of which become exposed upon its conformational activation[25]. We demonstrated that these cryptic epitopes are already accessible for recognition in patient samples from individuals with acute and subclinical iTTP[25, 33]. Our structural model readily explains these observations: in the compact latent state, distal modules such as TSP7 and TSP8 are bound at the MP domain, while the Linker bridges across MP, Dis, and Cys-rich exosite regions acting as a pseudosubstrate. This tight wrapping buries the corresponding surfaces and prevents antibody recognition. Once the distal domains are released during conformational activation, these surfaces become exposed, rendering them accessible to antibody binding.

To illustrate this conformational activation and inactivation process in our ADAMTS13-Del3To6 model, we performed a forced unfolding simulation (Sim6) separating the terminal CUB1-2 domains from the MDTCS core. The resultant force–distance profile is recorded in Figure 5. During unraveling, the MDTCS domains underwent a characteristic rotation that progressively released the distal domains from their surface in an unrolling motion, accompanied by an ordered sequence of rupture peaks in the force–distance trace. Although a single pulling trajectory cannot provide quantitative estimates of binding energies or the total mechanical work of unfolding, it nevertheless offers qualitative insights into the stability hierarchy of distal domain contacts. As may be expected, the two CUB domains exhibited the largest rupture forces, while the remaining distal domains disengaged more readily. Following the initial rupture, the subsequent domains are peeled away cooperatively, resulting in the global exposure of the MDTCS surface, explaining why in experiments with activating mAbs, and in plasma samples from iTTP patients, all cryptic epitopes become accessible simultaneously, regardless of their precise location (MP, Dis, TSP7, TSP8, Spacer, or CUB1)[25]. Conformational activation of ADAMTS13 appears to remodel the protein as a whole rather than selectively exposing individual epitopes.

**Figure 5:**
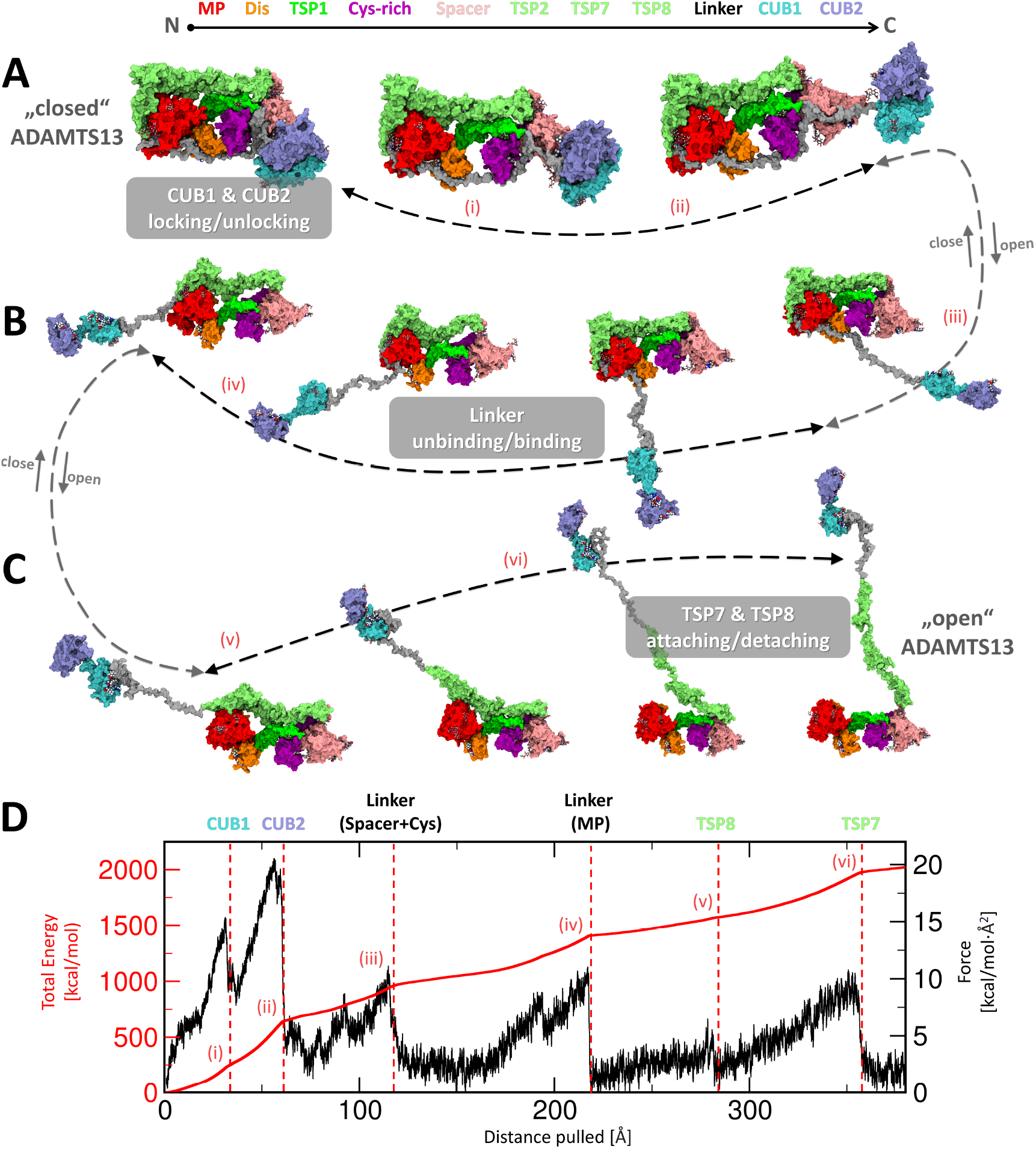
Conformational activation and inactivation mechanisms during global latency of ADAMTS13. Domain colors and sequential order are indicated at the top. **(A)** (left) Our final closed model of ADAMTS13-Del3To6 shows both terminal CUB domains attached to Spacer, the Linker region covers the substrate-binding sites on the MP, Dis, and Cys-rich domains while TSP7 and TSP8 bound near the MP domain. Pathway for conformational activation/inactivation illustrated with arrows. A movie illustrating this process intuitively is included in the provided Zenodo archive. During activation, the CUB1 and CUB2 domains detach from the Spacer module. Roman numerals (i-vi) highlight major rupture events observed during forced unfolding simulation (*Sim6*). **(B)** The Linker region further disengages, exposing substrate-binding exosites on the Cys-rich, Dis, and MP domains. **(C)** The TSP8 and TSP7 domains detach from the MP domain, completing the conformational activation of ADAMTS13. **(D)** Exemplary force and accumulated energy profiles derived from the forced unfolding simulation. (Structural snapshots relaxed with MD for natural appearance)

Interpreting data from *Zhu et al*.[21] on the proteolytic activity of ADAMTS13 constructs, lacking different distal domains (TSP7 through CUB2), in the light of our structural framework, suggests that conformational autoinhibition consists of two distinct stages, each contributing incrementally to the overall reduction in activity. Some of their truncated constructs displayed only partial inhibition, indicating that domain removal disrupts specific steps in the regulatory mechanism. In the first stage, loss of interactions involving TSP7 and TSP8 near the MP domain relieves approximately half of the autoinhibition, resulting in a two-fold increase in enzymatic activity. The second stage corresponds to the disruption of CUB1–2 domain binding to the Spacer module, producing an additional two-fold enhancement. Together, these transitions yield a roughly four-fold increase in activity for constructs unable to autoinhibit compared to the fully latent wild-type enzyme. ADAMTS13 may, hence, exist in a dynamic conformational equilibrium encompassing fully inhibited, partially active, and fully active states, providing a structural rationale for the low basal activity consistently detected in plasma, likely reflecting a minor subpopulation of molecules that transiently sample the open, active conformation even in the absence of external activators. Moreover, this framework explains the mechanism of conformational activation by mAbs: activating mAbs bind to epitopes transiently exposed in partially open states, stabilizing these intermediates and preventing their refolding into the autoinhibited form. Consequently, a progressively larger fraction of ADAMTS13 becomes trapped in the catalytically competent state, shifting the equilibrium toward activation.

Intriguingly, cryptic epitopes have been identified across most domains implicated in the conformational closure of ADAMTS13. In contrast, no cryptic epitopes were detected within the TSP3–TSP6 domains, which are dispensable for ADAMTS13 regulation. In this configuration, the distal domain chain is nearly fully extended, leaving no conformational slack to permit alternative wrapping modes. Consistent with this interpretation, activity data reported by Zhu et al. [21] indicate that constructs lacking additional distal domains display only partial autoinhibitory capacity. These constructs likely obstructed either the interaction interface near the MP domain or that near the Spacer domain, but obviously, not both simultaneously.

The compact wrapping of the distal domains acts as a molecular zipper that not only enforces autoinhibition but also enables its reversal. The Linker is of particular significance as it spans the MP, Dis, and the Cys-rich domains, thereby facilitating the CUB1–2 domains in proximity to the Spacer module. Only in this bound arrangement can CUB1–2 reliably re-engage with the Spacer module during the transition back into the latent state. Without such a molecular guide, reassociation of CUB1–2 would rely solely on diffusion over distances of several 100 Å, an event that is highly improbable in the crowded plasma environment. Thus, the zipper-like interplay of the Linker and distal domains provides a structural rationale for how ADAMTS13 can undergo repeated cycles of inactivation and activation.

### Conformational Responses to Chemical and pH Triggers

We explored conformational differences in plasma ADAMTS13 at physiological or sub-physiological pH. Hereto, binding to each of our cryptic epitope-recognizing mAbs was assessed in an in-house enzyme-linked immunosorbent assay (ELISA). In line with previous studies, almost no binding was observed for closed plasma ADAMTS13 under physiological conditions, suggesting that each cryptic epitope remained inaccessible (Figure 6, A, filled bars)[25]. Interestingly, sub-physiological pH conditions, that neutralize negatively charged residues and protonate His residues, open plasma ADAMTS13 and all cryptic epitopes became accessible, enabling distinct mAb binding (Figure 6, A, open bars). Hence, our findings indicate that ADAMTS13’s conformational changes are indeed pH sensitive, which fits the non-covalent nature of our simulated binding contacts. We and others have repeatedly shown that ADAMTS13 is allosterically activated at sub-physiological pH by an increase in its size and activity[34, 17, 35, 36, 24]. We hereby also associate this effect with the exposure of cryptic epitopes across multiple ADAMTS13 domains. This effect is consistent with our current findings, in which autoinhibition appears to rely heavily on networks of salt bridges and electrostatic contacts that serve to stabilize the compact domain arrangement. Acidic conditions would weaken or disrupt these interactions, in particular at interfaces where CUB1–2 bind to Spacer and where TSP7 and TSP8 are attached to the MP domain, facilitating their release exosite exposure.

**Figure 6:**
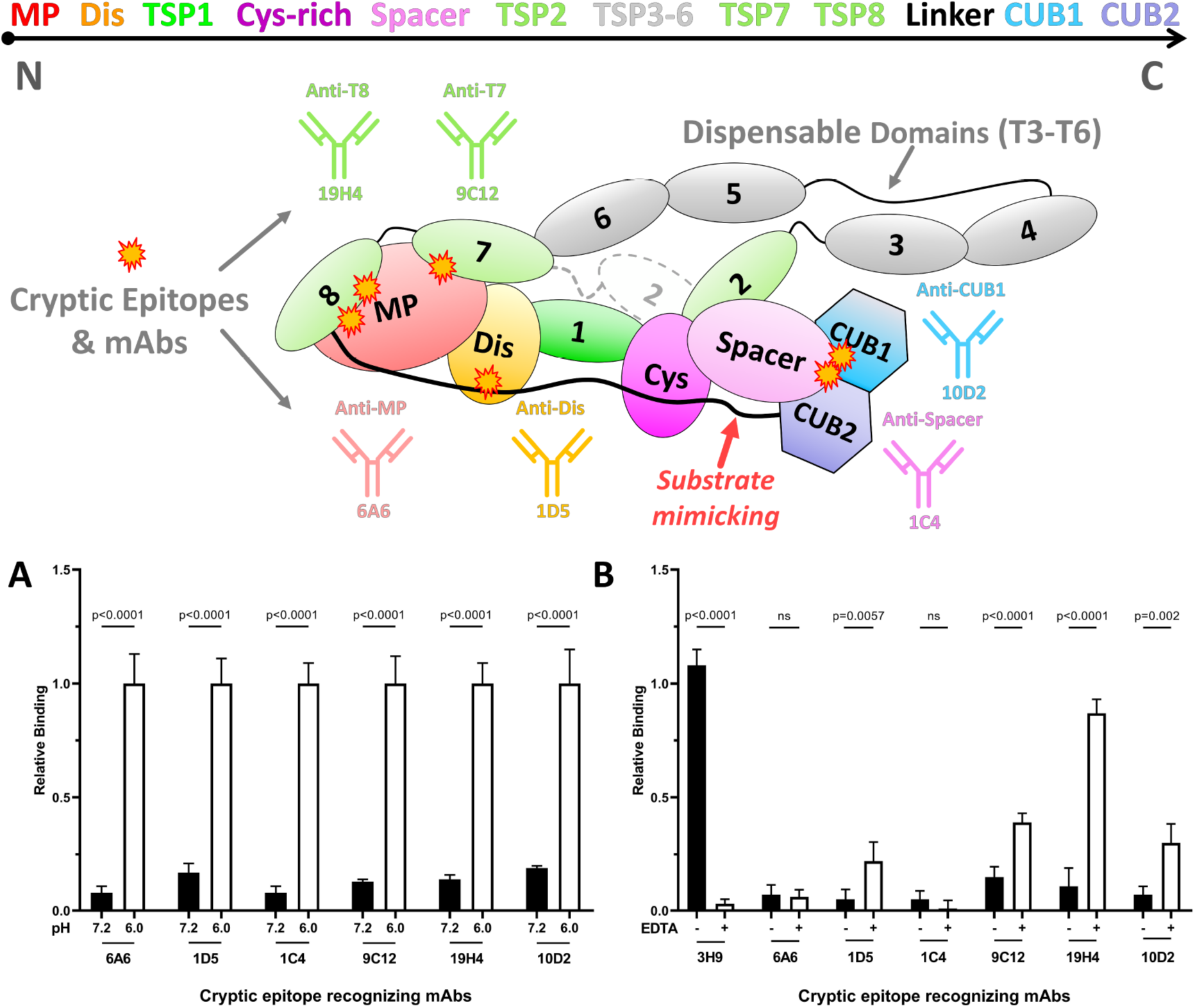
Conformation testing of ADAMTS13 under different conditions using ELISA.**(top)** Schematic depiction of updated model on conformationally latent ADAMTS13. Domain colors and sequential order indicated at the top. The CUB domains rests at Spacer. Other distal domains are wrapped around the substrate binding domains and shield off various cryptic epitopes for which mAbs were recently identified. The TSP3-6 repeats are dispensable. The Linker is a pseudosubstrate, acting like VWF-A2, binding to substrate exosites of MP, Dis, and Cys-rich modules. TSP7 and TSP8 rest at the MP domain. **(A)** Closed plasma ADAMTS13 incubated at pH 7.2 (filled bars) or pH 6.0 (open bars). In ELISA, binding of mAbs 6A6, 1D5, 1C4, 9C12, 19H4 and 10D2, directed against the MP, Dis, Spacer, TSP7, TSP8 and CUB1 domains, was assessed. All mAbs potently bind to open, but not to closed ADAMTS13. **(B)** Closed plasma ADAMTS13 was incubated in absence (filled bars) or presence (open bars) of EDTA. In ELISA, binding of mAbs was assessed as prevously. Chelation of Zn^2+^ and Ca^2+^ ions destabilized the structure of the MP domain resulting in abolished binding for mAbs 3H9 and 6A6, leading to partial unwrapping. (relative binding (mean ± one standard deviation) to open ADAMTS13 mediated by mAb 18H10. Significance tested using two-way ANOVA including multiple comparison with Bonferroni correction. All experiments were performed in triplicates (n=3))

To further investigate the aspects of our model around the direct inhibition of the MP domain, we aimed to locally alter the MP domain conformation without inducing extensive structural rearrangements as seen for mAb binding[28]. A Zn^2+^ and three Ca^2+^ ions are known to stabilize ADAMTS13’s MP domain conformation and warrant ADAMTS13’s functionality[8]. Hence, we incubated closed plasma ADAMTS13 in the absence or presence of 10 mM ethylenediaminetetraacetic acid (EDTA) to chelate both Zn^2+^ and Ca^2+^ ions. As expected, only the non-cryptic mAb 3H9 showed binding to plasma ADAMTS13 in the absence of EDTA, and all cryptic epitopes remained inaccessible (Figure 6, B, filled bars). With ions only being present in the MP domain, EDTA incubation was expected to only locally alter the MP domain structure. Indeed, chelation of Zn^2+^ and Ca^2+^ ions revealed abolished binding of the anti-MP mAbs 6A6 and 3H9 (Figure 6, B, open bars), indicating structural changes to the MP domain. Interestingly, ion chelation also induced potent binding of mAb 19H4, indicating the accessibility of the cryptic TSP8 epitope as well. Hence, chemical destabilization of the MP domain also detached the TSP8 domain interaction proposed in our new model for closed ADAMTS13. Although to a lesser extent, mAbs 1D5 and 9C12 were able to bind following ion chelation. This suggests that the detachment of the TSP8 from the MP domain likely also uncouples TSP7 and the Linker in this region, allowing mAb binding at the cryptic Dis epitope, too. Furthermore, EDTA-mediated ADAMTS13 opening also revealed modest binding of anti-CUB1 mAb 10D2, which likely reflects the uncoupling of the Spacer-CUB1 interaction. Unfortunately, we were unable to further corroborate this finding as no mAb 1C4 binding was observed due its abolished detection by the biotinylated anti-MP mAb 3H9 (Figure 6, B, open bars).

Taken together, ADAMTS13 remains closed through non-covalent interactions, as cryptic epitope exposure was found to be sensitive to pH changes. This finding fits the hydrogen–deuterium exchange mass spectrometry (HDX-MS) reports on peptides from various ADAMTS13 domains that show deuterium uptake alterations when incubated at different pH conditions. HDX-MS experiments by Pillai *et al*.[24] revealed significant protection differences between full-length ADAMTS13 and truncated MDTCS variants, while we have shown altered deuterium uptake within the terminal CUB domains in the absence and presence of activating mAbs[28]. Regions within the MP, Cys-rich, Spacer, and both CUB domains displayed altered deuterium uptake compared to the inactive protein, indicating that these segments are shielded from solvent in the latent conformation. Our ADAMTS13-Del3To6 model provides a direct structural rationale for these observations (Figure S18). Chelation of Zn^2+^ and Ca^2+^ ions not only destabilized the MP domain structure but also enabled potent or modest mAb binding to cryptic epitopes. These EDTA-mediated structural changes are in line with the previously reported increase in hydrodynamic diameter measured by dynamic light scattering, and the reduced *α*-helix and *β*-sheet content found by FTIR spectroscopy[37]. Furthermore, we characterized the molecular process of enzyme unfolding following demetallation since the potent mAb 19H4 binding reflects the uncoupling of the TSP8 domain from the MP domain as proposed in our model. Modest binding of mAb 9C12 also suggests the detachment of the TSP7 domain from MP. However, it is unclear if the cryptic Dis domain epitope is blocked by TSP7 or the Linker. Although 1C4 epitope exposure could not be detected, modest binding of mAb 10D2 also suggested the uncoupling of the Spacer and CUB domains, again reflecting the same collective release of distal domains discussed before.

## Conclusion

Our study establishes the first cohesive structural model of ADAMTS13 autoinhibition that integrates and mechanistically explains all major experimental observations. In the globally latent state, TSP7, TSP8, the Linker (L3), and the CUB1–2 domains form a compact assembly that wraps around the MDTCS core, sterically blocking the catalytic cleft and masking key VWF-binding exosites. A central insight is that the Linker acts as a pseudosubstrate resembling VWF-A2, binding across MP, Dis, and Cys-rich exosites while guiding CUB1–2 to Spacer. This “molecular zipper” stabilizes the autoinhibited architecture and enables efficient, reversible transitions between closed and open states. The compactness of this arrangement explains why ADAMTS13-Del3To6 is the shortest construct capable of full autoinhibition and aligns with the extended hairpin organization suggested by SAXS data for full-length enzyme. Our simulations further reveal refined, distinct binding modes for CUB1 and CUB2 that require release from the crystallographic tandem configuration, clarifying their differing inhibitory contributions.

Pulling simulations show that activation proceeds through an ordered release of distal domains. Initial Spacer–CUB disengagement, followed by Linker detachment and unwrapping of TSP7 and TSP8 lead to global exposure of cryptic epitopes in agreement with antibody-binding. This framework resolves long-range regulatory effects by demonstrating how Spacer–CUB interactions indirectly shape MP accessibility and activity.

Environmental perturbations support this model: mildly acidic pH or chelation of Zn^2+^ and Ca^2+^ ions weakens electrostatic interfaces within MP and among distal domains, inducing partial opening and exposing otherwise hidden epitopes.

Together, these findings provide a unified structural and mechanistic explanation for ADAMTS13 autoinhibition and its relief. The model rationalizes longstanding observations concerning cryptic epitope exposure, distal-domain contributions, and the remote influence of Spacer–CUB interactions on MP activity. It also establishes a foundation for therapeutic strategies aimed at modulating ADAMTS13 function by stabilizing or disrupting specific domain–domain interfaces, with direct relevance for treating TTP and other disorders of the ADAMTS13–VWF axis

## Data availability statement

The data that support the findings of this study are available within the article and its supplementary materials. The PDB files of the major models, full trajectories for all simulations and resulting clusters are openly available in Zenodo at: https://doi.org/10.5281/zenodo.17181850. The data archive also includes a video showing the opening of our ADAMTS13-Del3To6 model during the pulling simulation, illustrating the mechanism intuitively.

## Materials and Methods

Full methodological details are provided in the Supplementary Information. In brief, we combined extensive atomistic simulations and biochemical experiments to construct and test a structural model of ADAMTS13 global latency. A shortened human ADAMTS13 construct lacking TSP3–TSP6 (ADAMTS13-Del3To6; UniProt Q76LX8) was built using AlphaFold2 and available crystal structures as templates, including the MDTCS core (PDB ID 6QIG) and the CUB1–2 tandem (PDB ID 7B01).

To explore distal-domain binding and autoinhibition, we performed extensive molecular dynamics simulations with NAMD 2.14 and the CHARMM36 force field in explicit solvent. Enhanced sampling was achieved with our TIGER2h_*P E*_ replica-exchange protocol, which combines explicit-solvent simulations with implicit-solvent energies for exchange decisions. Simulations were used to probe TSP7/TSP8 binding to the MP domain, substrate-like binding of the Linker region to MDTCS, and wrapping of distal domains when CUB1–2 are anchored to Spacer. Clustering analyses were used to derive representative structures. Constant-velocity pulling was carried out to visualize the coordinated unwrapping and exposure of cryptic epitopes in the final model. Experimentally, conformational states of plasma-derived ADAMTS13 were examined using ELISA with monoclonal antibodies recognizing cryptic or non-cryptic epitopes across the MP, Dis, Spacer, TSP7, TSP8, and CUB1 domains. Plasma ADAMTS13 was incubated at physiological or mildly acidic pH, or in the presence of 10 mM EDTA to chelate Zn^2+^ and Ca^2+^ ions. Antibody binding was normalized to an open-state reference antibody, and statistical comparisons were performed using two-way ANOVA with Bonferroni correction.

## Acknowledgment

This work was supported by the North-German Supercomputing Alliance (HLRN), grant number mvb00016, and by the European Research Council (ERC), Starting Grant “PredicTOOL” (637877) to M.D.

## Competing interests

There are no competing interests to declare.

## CRediT authorship contribution statement

**Norman Geist:** Conceptualization, Methodology, Software, Investigation, Data Curation, Writing - Original Draft, Visualization. **Quintijn Bonnez:** Conceptualization, Methodology, Investigation, Data Curation, Formal analysis, Writing Original Draft, Visualization. **Karen Vanhoorelbeke:** Resources, Writing - Review & Editing, Supervision, Funding acquisition, Validation. **Mihaela Delcea:** Resources, Writing - Review & Editing, Supervision, Funding acquisition, Validation.

## Notes

https://doi.org/10.5281/zenodo.17181850

### Competing Interest Statement

The authors have declared no competing interest.

