## Extended Methods and Supplementary Figures PDF for "Wrapping it Up: Structural Basis of ADAMTS13 Global Latency"

### Appendix

#### Extended Materials and Methods

##### Simulation settings

Simulations were implemented with NAMD 2.14[38], its TCL programming language API, and the COLVARS module[39]. Full periodic boundary conditions are applied, and long-range electrostatics are considered with the smooth-particle-mesh-ewald[40] in a pseudo-NVT ensemble[41]. Short-range interaction cutoffs were set to 1.0 nm with a switching function of 0.1 nm. Pressures and temperatures were maintained by a Langevin piston barostat at 100 fs period and 200 fs decay time and a Langevin thermostat with a 1 $ps^{-1}$ damping coefficient. Parametrization of proteins and glycans was done with CHARMM-GUI and the CHARMM36 forcefield[42]. Solvation boxes with the TIP3P[43] water model and neutralizing sodium or chloride ions were added with VMD 1.9.4[44], while hydrogen mass repartitioning[45] (HMR) was implemented with PSFGEN, allowing an increased timestep of 4 fs.

##### Replica-exchange simulations (TIGER2h$_{PE}$)

Multiple replicates of the simulation with full explicit solvation proceed at different temperatures as in conventional T-REMD[46], but exchange decisions are based on energies obtained from implicit solvent models and hence, TIGER2h$_{PE}$ can rely on a greatly reduced number of replicas. Meanwhile, accuracy is strongly improved compared to pure implicit solvent simulations. Exchange attempts are carried out between unique pairs of randomly selected replicas after the cooling phase according to an updated TIGER2h$^{PE}$ scheme, with the subsequent sorting by potential energies discarded[26]. After temperatures have been swapped, reheating and sampling follow for another cycle. This procedure repeats until a sufficiently converged state ensemble is collected at the baseline temperature. Due to the presence of multiple solutes, no explicit solvation shell was considered. Implicit solvent energies used for exchange decisions were evaluated by the GB$_{OBCII}$[47] model using OpenMM[48] and periodic boundary conditions. Non-bonded cutoffs were set to half the cell size. The sampling and cooling phases were set to 32 ps and 8 ps for all simulations, except Sim2, which used a sampling phase of 16 ps. The temperature ladder spanned from 300 K up to 450 K. Each simulation started with an energy minimization equivalent to two replica-exchange cycles' worth of steps.

##### Simulated systems

The shortened ADAMTS13 model lacking the TSP repeats three to six was generated based on the sequence information from Uniprot[49] under accession code Q76LX8. AF2 was employed to generate a structural model with high confidence from the core MDTCS and TSP2 domains (80-745), then continuing with TSP7, TSP8, Linker, and the terminal CUB1-2 domains (1016-1410)(Figure S1). The core MDTCS domains were subsequently replaced by the PDB ID 6QIG chain C (79-682), including the zinc ion in the active center, three calcium ions in the MP domain, and seven glycan chains distributed across the surface of MDTCS. Missing amino acids in the Cys-rich domain (454-468) were partly included from PDB ID 3GHM (452-458 and 466-469). The region 459-465 was added as a random coil. The present Q225E mutation in 6QIG was reverted with CHARMM-GUI[50] during parameterization. One glycan chain was imported at CUB1 from PDB ID 7B01 via structural alignment.

###### *Sim1*

A model truncated after TSP8 was solvated in a cubic box with 200 Å side lengths to sample the conformations of TSP7 and TSP8 under natural constraints.

For this and all other simulations, the accessible conformational space of the ADAMTS13-MDTCS domains was restricted by two separate RMSD flat-bottom restraints (backbone): one spanning MP through TSP1 and the other spanning TSP1 through Spacer. Because the MP–Dis and Cys-rich–Spacer domain pairs share extensive interfaces, applying separate structural restraints to each domain would overly limit conformational flexibility. The restraints limited backbone RMSD deviations from the initial structure to 15 Å with a force constant of 5 kcal $mol^{-1}$. Additional restraints were applied to each TSP2–TSP8 repeat individually, with a threshold of 2.5 Å at the same force constant. Finally, ADAMTS13-MP residues coordinating calcium or zinc ions were maintained in their crystal structure geometry using additional harmonic bonds with a force constant of 1 kcal $mol^{-1}$.

To refine the cluster analysis of the resulting state ensemble, we filtered conformational states to retain only structures in which TSP7 or TSP8 interacted with the MP or Dis domains (39 % of structures) for subsequent RMSD cluster analysis (Figure S2). From the five extracted structural clusters, three displayed similar coiled-up binding poses of TSP7 and TSP8 at the MP domain: clusters 0 (44.66 %), 1 (15.74 %), and 3 (4.15 %). These can be interpreted as artifacts of the truncation after TSP8. However, two binding poses of TSP8 attached to the MP domain were returned: clusters 2 (9.09 %) and 4 (3.50 %), which were suitable for the connection with the Linker within the active site cleft.

**Table S1**
**Overview over TIGER2h$_{PE}$ and final pulling simulations**. Columns three to seven denote the number of replicas (#R), total simulation time (quenching excluded) at the baseline temperature in nanoseconds, total number of states, average temperature change per exchange and average exchange rate.

| Nr. | System Components | #R | Tbase [ns] | Nstates | DT/X [K] | P(X) |
|---|---|---|---|---|---|---|
| 1 | ADAMTS13-MDTCS-T278 | 8 | 1430 | 46559 | 8.84 | 0.21 |
| 2 | ADAMTS13-MDTCS + Linker | 16 | 973 | 60655 | 11.03 | 0.29 |
| 3 | ADAMTS13-Del3To6 | 16 | 647 | 20136 | 8.56 | 0.24 |
| 4 | ADAMTS13-Del3To6 | 16 | 598 | 18692 | 9.45 | 0.26 |
| 5 | ADAMTS13-Del3To6 | 16 | 1201 | 37448 | 7.97 | 0.23 |
| 6 | ADAMTS13-Del3To6 (Pulling) | - | - | - | - | - |

##### *Sim2*

A model truncated after the Spacer domain is employed, while the linear Linker model was generated using the AmberTools24[51] package and parameterized with CHARMM-GUI. The Linker was positioned as a linear peptide adjacent to the MDTCS domains in the correct orientation, with the N-terminus placed near the MP domain and the C-terminus directed toward the Spacer module. The system was solvated in a rectangular cell measuring 210 Å × 80 Å × 80 Å.
In classical force fields, the absence of charge polarization effects can cause unnaturally strong interactions between the carboxyl groups of glutamic and aspartic acid residues and the zinc ion in the ADAMTS13-MP active site. To prevent this artifact, protective flat-bottom restraints were applied, repelling these side chains when the distance fell below 4.5 Å, with a force constant of 50 kcal mol$^{-1}$. This was applied to MP residues E225 and D252, as well as Linker residues E1142 and E1143.
To avoid unproductive exploration of Linker binding outside the active site cleft or positional drift toward the termini, a guiding restraint was introduced to improve sampling efficiency. In our previous work, we enforced binding between the carbonyl oxygen of the catalytic Y1605 residue of VWF-A2 and the MP zinc ion. Since no such catalytic interaction is expected for the Linker, we adopted a different strategy: a two-sided RMSD flat-bottom restraint on the backbone between MP binding pocket residues (L147–T148, L151, L185–P186, N189, V195–L198, W206, H234) which previously engaged VWF-A2 residues (P1601–V1604) and the homologous Linker segment (P1136–V1139) identified in our proteolytic binding Model1. This assumed that the Linker would bind the MP domain at the same location as VWF-A2.
To further promote Linker engagement with the Dis and Cys-rich domains, the centers of their proposed exosites (Dis: R349-L350, Cys-rich: A472-V474) were kept near their corresponding regions on the Linker (Dis: 1141-1149, Cys-rich: P1159-L1168) by sequence similarity, using soft flat-bottom restraints between the centers of mass of atom groups, identical to our VWF-A2 simulations[14]. Only when the upper distance limit of 25 Å was exceeded could these Linker regions not detach further from their proposed MDTCS interaction sites. Finally, to mimic the omnipresent tensile force acting on VWF-A2 under physiological conditions, a constant pulling force of 1 pN was applied at both termini of the Linker during sampling, too. This also served to prevent coiling of the Linker.
The structural ensemble was subsequently filtered to contain only structures with contacts between the Linker and the Dis exosites (R349–L350) and both Cys-rich exosites, the one from our previous work (I495–K497) and the earlier proposed (A472–V473). As the transition between the MP and Dis binding portions of the Linker is less clear compared to VWF-A2 due to the strong interaction of the **EE** motif with the MP domain, no further filters were applied to these regions. However, the Cys-rich binding position of the Linker, extended by the **PR** motif (L1159–R1176), was filtered for contacts with the whole Cys-rich module and subjected to RMSD cluster analysis. A total of 12 % of frames remained. After alignment of the MDTCS core domains, the RMSD clusters of the corresponding binding regions on the Linker for the MP, Dis, and Cys-rich modules, analogous to the VWF-A2 domain, were searched (Figure 2, A). The first three clusters are as follows: 0 (48.83 %), 1 (16.27 %), and 2 (3.33 %), yielding a tight binding structure of the Linker region to the MDTCS domains, with only marginal variations. The remaining clusters displayed less structured binding poses near the Cys-rich module or a large offset from the Dis exosite, only ranging within 2.17-1.87 %.

##### *Sim3-5*

The first simulation of the full ADAMTS13-Del3To6 model (Sim3) was initiated by combining the results from Sim1 and Sim2. The TSP2–TSP8 structure from Sim1, with TSP8 attached to the MP domain, was merged with the binding pose for the Linker from Sim2, ensuring a continuous connection to TSP8 and also allowing the two terminal CUB domains to be appended. Therefore, a Linker binding state was selected that showed interaction with the Cys-rich exosite (I495–K497), but with the remaining Linker towards Spacer unbound. The CUB1-2 domains were pre-aligned according to the proposed binding pose by Kim *et al.* [9], but positioned at a fair distance from the Spacer module. In addition to the analogous structural restraints on protein modules already mentioned for the previous simulations, the CUB1-2 domains were also protected by a joint RMSD flat-bottom restraint, keeping them close to their crystal structure state (PDB ID: 7B01). The system was solvated in a cubic cell with side lengths of 150 Å. Sampling was restarted from a representative state of Sim3, showing further attachment of TSP7 to the MP-Dis interface (Sim4). The final sampling (Sim5) was based on a representative state from Sim4, but with the CUB domains released from their fused configuration. Instead, RMSD flat-bottom restraints were applied individually to each CUB domain. For CUB1, the restraint was additionally used as a positional guide, based on the best Spacer–CUB1 binding model predicted by AF2, while CUB2 remained free to discover its binding pose. The system was then solvated in a cubic box with side lengths of 150 Å. To obtain the final model from Sim5, we filtered the structural ensemble for contacts between the Spacer module and the critical CUB1 and CUB2 residues reported by Kim *et al.*[9]. This resulted in retention of only 6.2 % of structures, which were then subjected to RMSD-based clustering using the C$\alpha$ atoms of both CUB domains after alignment of the Spacer domain. This yielded three major clusters: cluster0 (51.33 %), cluster1 (24.4 %), and cluster2 (19.9 %). Clusters 0 and 1 displayed highly similar structures, differing only in slight displacements of the CUB domains but sharing the same binding interfaces with Spacer. All key residues of CUB1 and CUB2 engage directly with the Spacer domain. The interface also includes the well-established Spacer residues R660, Y661, and Y665, which are critical for VWF-A2 binding and mAb recognition. In addition, due to the fact that the remaining distal domains in cluster0 are also tightly engaged with the MDTCS core, yielding a coherent and stable arrangement, cluster0 of Sim5 is designated as the final conformationally inactive model of ADAMTS13.

##### *Sim6*

An exemplary pulling simulation was carried out to probe the new conformationally inactive ADAMTS13-Del3To6 model and to visualize the wrapping and unwrapping process step by step. A conventional MD simulation under implicit solvent conditions (GB$_{OBCII}$[47]) was performed, with constant-velocity pulling applied through the Colvars module to increase the distance between the centers of mass of the MDTCS domains and both CUB domains to more than 1000 Å over the course of 200 ns. The pulling rate was set to 5 Å ns$^{-1}$ with a spring constant of 1 kcal mol$^{-1}$. To preserve the overall secondary and tertiary structure during the pulling, the same RMSD flat-bottom restraints used in Sim5 were applied.

#### Sequence alignment

Sequence similarity between the VWF-A2 domain and the Linker region of ADAMTS13 was assessed using the UniProt Alignment tool[49]. The amino acid sequence of the human VWF-A2 domain (UniProt accession: P04275, residues 1498-1665) and the human ADAMTS13 sequence (UniProt accession: Q76LX8, residues 1132-1191) were used. Parameters were kept at default settings. For visualization and functional annotation of conserved regions, the TEXSHADE package in LaTeX was used.

#### Structural filters

The structure ensembles produced from sampling simulations were filtered using VMD and its TCL interface prior to performing cluster-based analysis. This filtering step removed any states lacking atomic contacts between two specified selections within a set threshold. Consequently, structures that did not meet the criteria for proposed exosites and interactions were excluded. Since no direct contact pairs are reliably available from experimental data, for exosites, the filtering process was carried out in two stages. First, structures were checked for contacts between the proposed exosite on one ADAMTS13 module and the entire surface of another module. Next, contacts were identified between the proposed region of the second module and the entire surface of the first ADAMTS13 domain. This approach ensures a comprehensive evaluation of the proposed regions critical for interactions between ADAMTS13 modules.

#### Clustering of binding modes

To identify favorable binding structures between ADAMTS13 components in our simulation, we utilized RMSD cluster analysis using VMD. After the alignment of the respective stationary component, e.g., MDTCS in Sim1, RMSD cluster analysis was carried out on the respective movable parts, e.g., the Linker in Sim1, always searching for five clusters with a 6 Å cutoff.

#### Binding affinities and contact statistics

Contact statistics in complexes were evaluated by a parallel in-house code that counts individual residue contacts between both proteins and has already been proven effective in earlier works[14]. A contact is characterized by atoms of any two residues between two selections coming closer than 2.5 Å. Thereby, persistent contacts and possibly important residues for the binding can be identified. The counts are normalized to the number of frames to represent the average fraction of contacts per residue throughout the simulation, and are used to interpret binding affinities. Affinity values greater than one indicate binding to several other residues simultaneously. On the one hand, the contact statistics can be displayed in a network-like diagram, which allows the evaluation of specific individual contacts. On the other hand, they can be averaged from the perspective of both proteins to identify important residues and surface areas in general. Increased specificity of contacts is seen when individual compounds in the network plot have high affinity. Low specificity, in contrast, is seen when there are many interactions of a residue with only low affinities, especially when these contacts occur alternately rather than simultaneously. In this case, this indicates multiple binding sites or a non-specific interaction. However, if multiple low-affinity contacts occur simultaneously, this is likely an indication of a residue that is bound in a specific binding pocket.

#### ADAMTS13 conformation testing in ELISA

To support our model for closed ADAMTS13, the exposure of cryptic epitopes within the MP, Dis, Spacer, TSP7, TSP8 and CUB1 domains of plasma ADAMTS13 was investigated under various pH or chelating conditions. Hereto, a previously described in-house ELISA assay was slightly modified[25]. All assays required plasma ADAMTS13 and, therefore, plasma from >20 healthy donors was pooled. All individual plasmas were obtained in accordance with the Declaration of Helsinki under local ethical committee approval (S62889 and S66725, UZ/KU Leuven, Belgium). All mAbs (3H9, 6A6, 1D5, 18H10, 1C4, 15D1, 9C12, 19H4, 10D2 and 17G2) were produced and purified as previously described[25]. In brief, (non-) cryptic epitope-recognizing mAbs were individually coated (5 µg/mL in 0.05 M (bi)carbonate buffer at pH 9.6). After blocking, serial dilutions of NHP plasma were prepared using 5 mM Bis-Tris buffer at pH 7.2 or pH 6.0. When exploring chelating conditions, NHP plasma was serially diluted using 0.3 % skimmed milk powder in PBS containing a final concentration of 10 mM EDTA. As an intra-assay reference and positive control for open ADAMTS13, NHP plasma was serially diluted using 0.3 % Milk/PBS containing 5 µg/mL of the anti-T1 mAb 18H10. To detect captured, open ADAMTS13, the biotinylated anti-Spacer mAb 15D1 (1.5 µg/mL in 0.3 % Milk/PBS) and HRP-labeled high-sensitivity streptavidin (1/10 000 in 0.3 % Milk/PBS; PierceTM, Waltham, USA) was used. As the cryptic epitope-recognizing mAb 1C4 already binds ADAMTS13's Spacer domain, the biotinylated antibody 3H9 was used for detection as previously described[25, 52]. Color reactions were induced by oxidation of Ortho-phenylenediamine and hydrogen peroxide and terminated after 10 minutes using 4 M sulfuric acid. Optical density (OD) was measured at 492 nm and relative binding was determined by normalizing OD values against those of the 18H10 intra-assay positive control[36].

**Supplementary Figures**

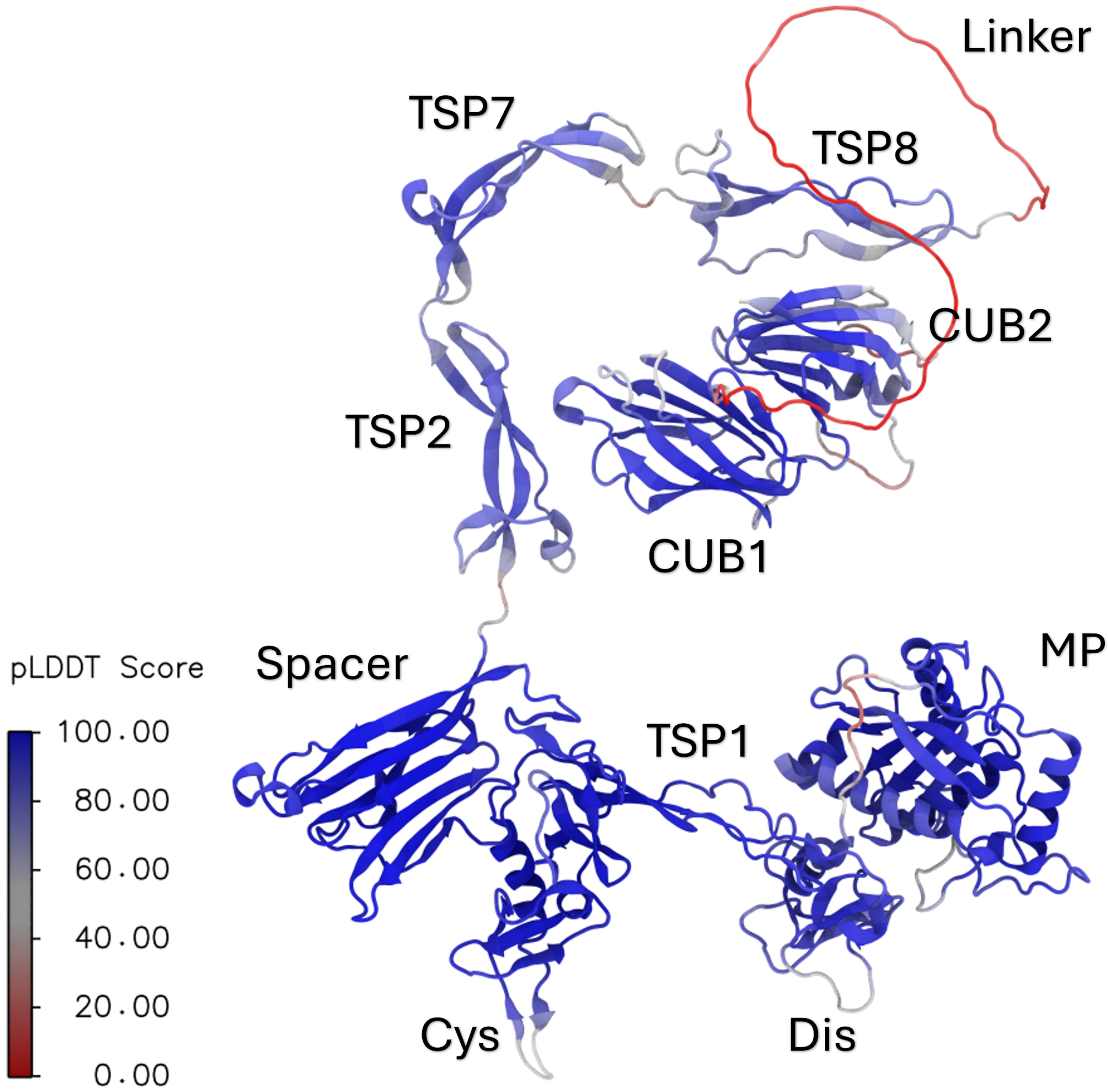


**Figure S1: ADAMTS13-Del3To6 model as obtained through AlphaFold2, colored by pLDDT score.**

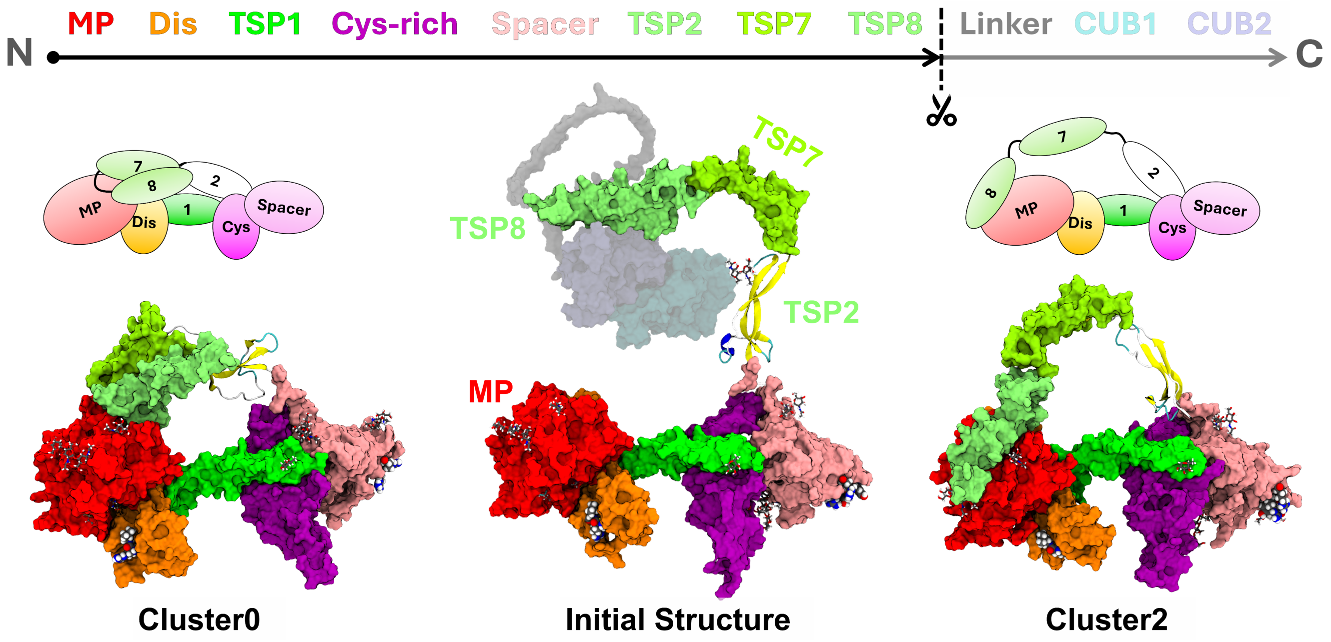


**Figure S2: Structural sampling of TSP7 and TSP8 in a ADAMTS13-Del3To6 model truncated after TSP8.** Results from (*Sim1*) illustrate the accessible positioning of TSP8 at the MP domain under natural constraints. (Top) The domain colors and their sequential order from N- to C-terminal are indicated. (Middle) Initial structure as predicted by AF2, with domains truncated after TSP8, visualized by faded colors. (Left) One major structural pose as obtained through RMSD cluster analysis of the state ensemble sampled by TIGER2h$_{PE}$. TSP7 and TSP8 are bound near the MP module in a coiled fashion. (Right) Different structural clusters show TSP8 bound to the MP domain in a way that allows connection with the Linker region when binding in a substrate-like fashion across the active site.

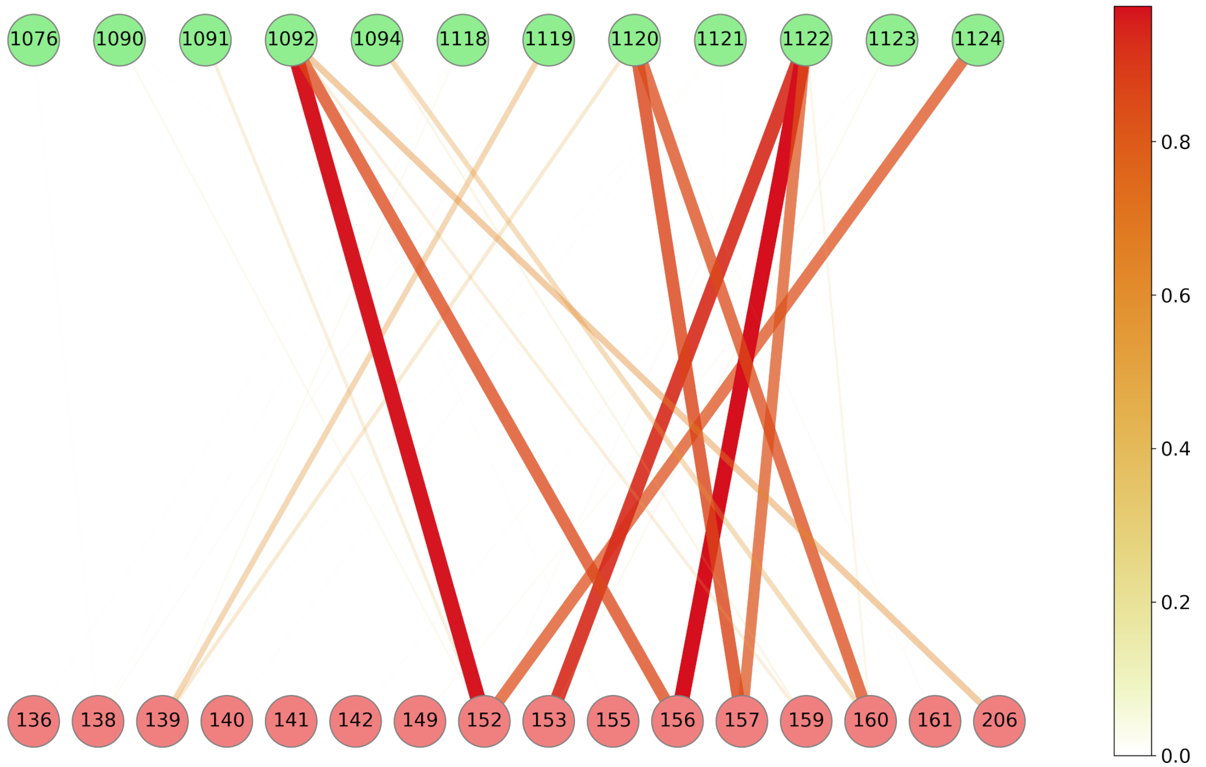


**Figure S3: Residue interaction network of MP domain (red) with TSP8 (green) for cluster2 frames from Sim1.** Contacts of affinities below 0.1 were cut off. The connection line thickness and transparency are quadratically weighted by the affinities and colored from red to white for high and low affinity values, respectively.

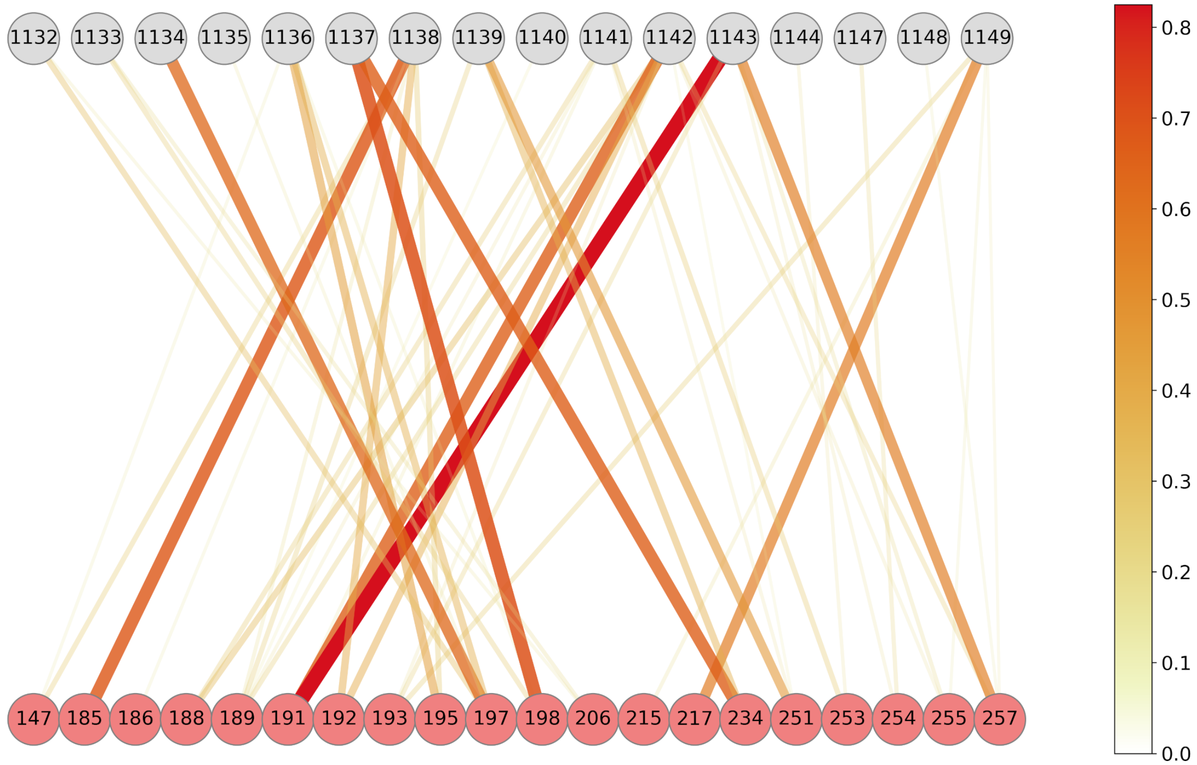


**Figure S4: Residue interaction network of MP domain (red) with Linker (gray) for cluster0 frames from Sim2.** Contacts of affinities below 0.1 were cut off. The connection line thickness and transparency are quadratically weighted by the affinities and colored from red to white for high and low affinity values, respectively.

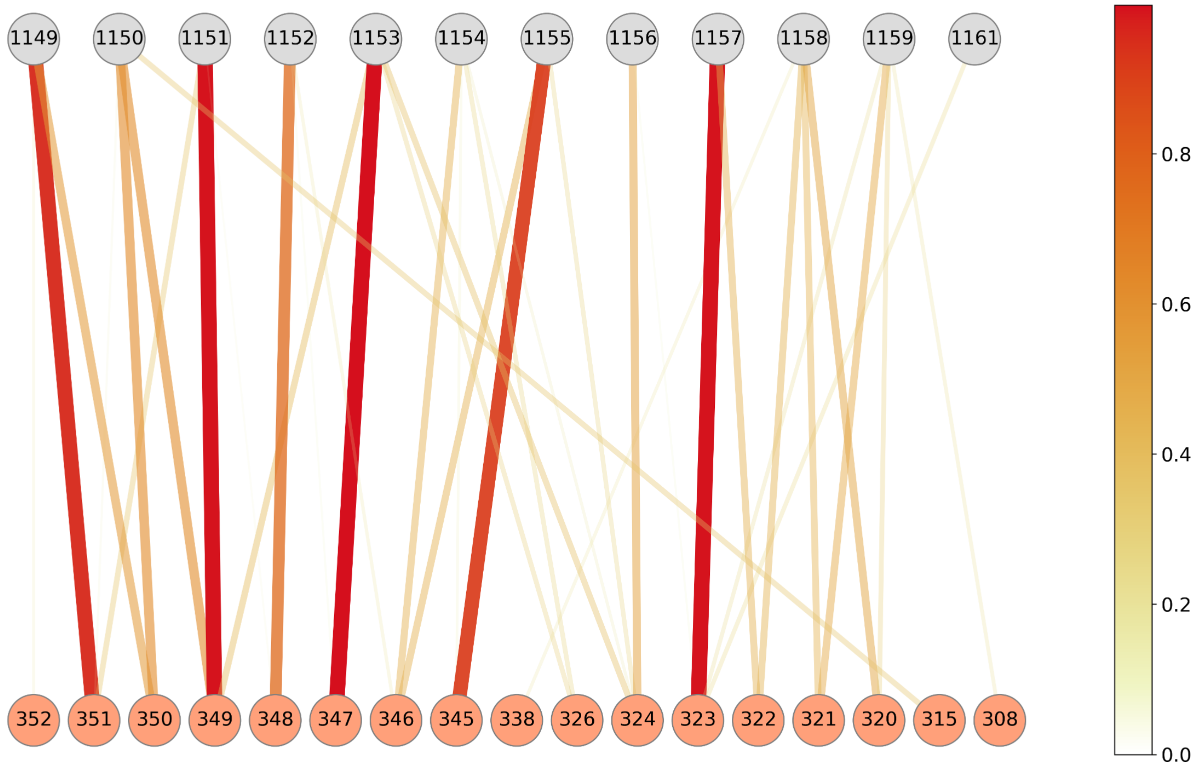


**Figure S5: Residue interaction network of Dis domain (orange) with Linker (gray) for cluster0 frames from Sim2.** Contacts of affinities below 0.1 were cut off. The connection line thickness and transparency are quadratically weighted by the affinities and colored from red to white for high and low affinity values, respectively.

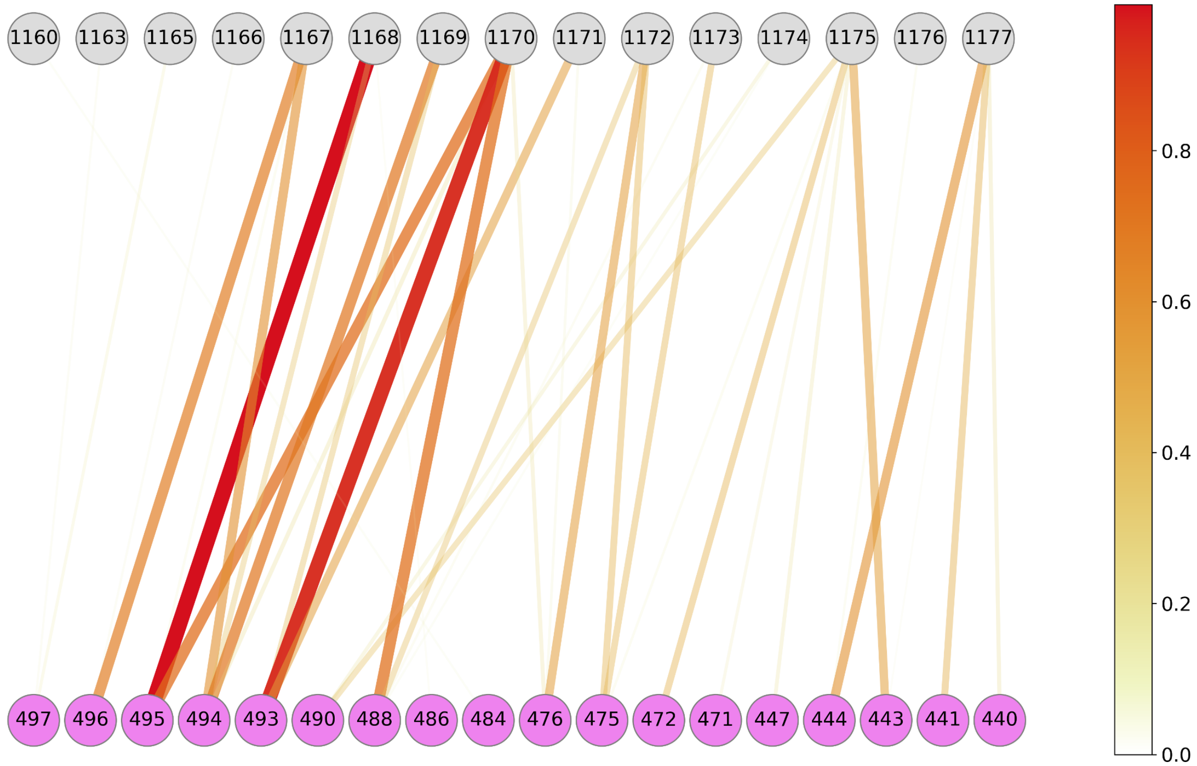


**Figure S6: Residue interaction network of Cys domain (purple) with Linker (gray) for cluster0 frames from Sim2.** Contacts of affinities below 0.1 were cut off. The connection line thickness and transparency are quadratically weighted by the affinities and colored from red to white for high and low affinity values, respectively.

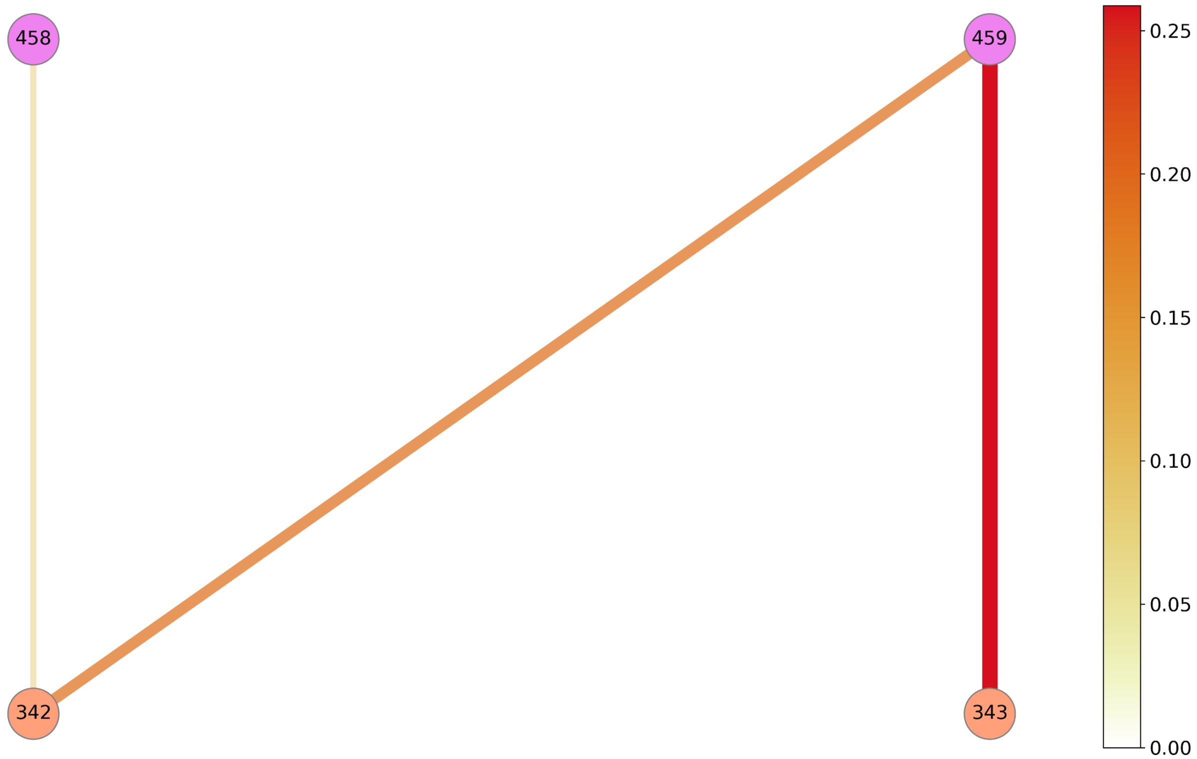


**Figure S7: Residue interaction network of Cys domain (purple) with Dis domain (orange) for cluster0 frames from Sim2.** Contacts of affinities below 0.1 were cut off. The connection line thickness and transparency are quadratically weighted by the affinities and colored from red to white for high and low affinity values, respectively.

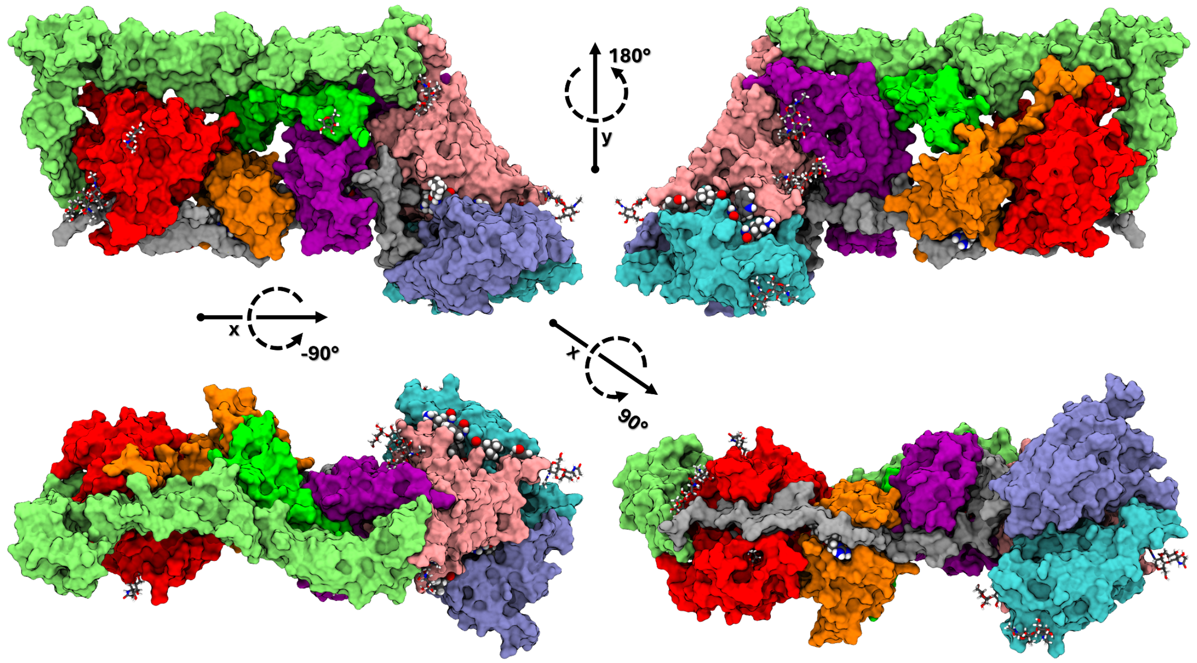


**Figure S8: Surface representation and different viewing angles for final conformationally inactive ADAMTS13-Del3To6 model as obtained through Sim5 (cluster0)**

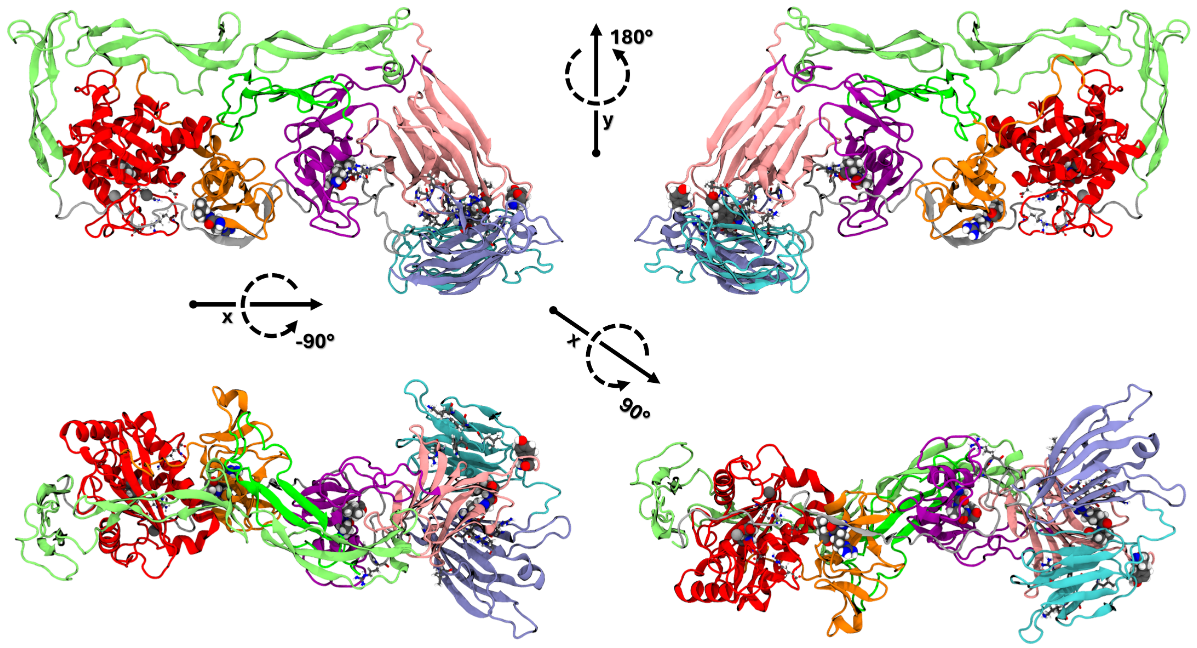


**Figure S9: Cartoon representation and different viewing angles for final conformationally inactive ADAMTS13-Del3To6 model as obtained through Sim5 (cluster0)**

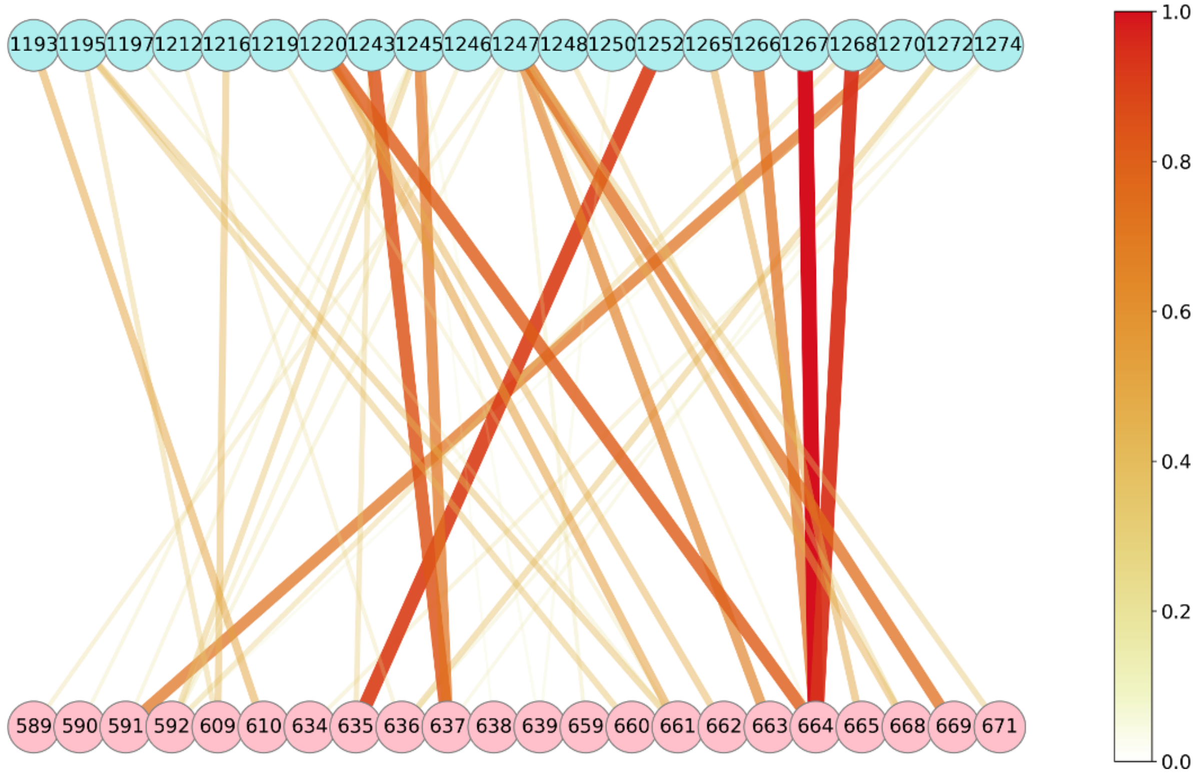


**Figure S10: Residue interaction network in final model of Spacer domain (pink) with CUB1 (cyan) for cluster0 frames from Sim5.** Contacts of affinities below 0.1 were cut off. The connection line thickness and transparency are quadratically weighted by the affinities and colored from red to white for high and low affinity values, respectively.

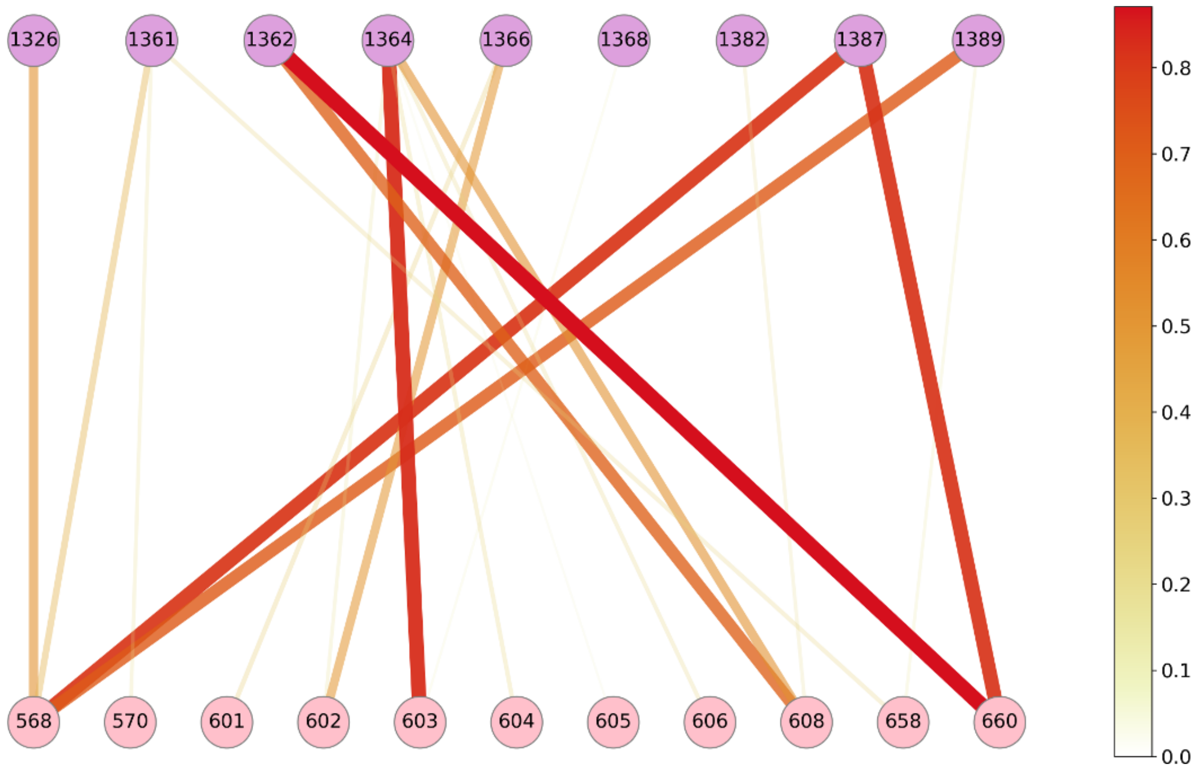


**Figure S11: Residue interaction network in final model of Spacer domain (pink) with CUB2 (plum) for cluster0 frames from Sim5.** Contacts of affinities below 0.1 were cut off. The connection line thickness and transparency are quadratically weighted by the affinities and colored from red to white for high and low affinity values, respectively.

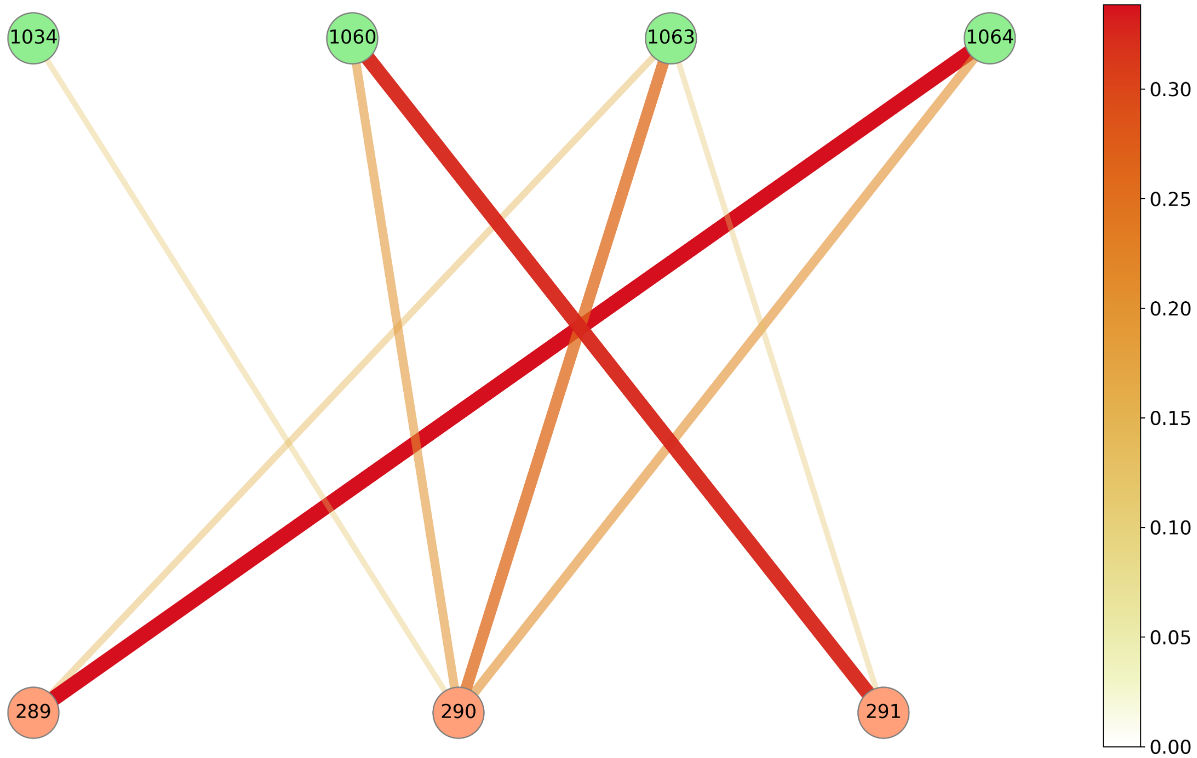


**Figure S12: Residue interaction network in final model of Dis domain (red) with TSP7 (green) for cluster0 frames from Sim5.** Contacts of affinities below 0.15 were cut off. The connection line thickness and transparency are quadratically weighted by the affinities and colored from red to white for high and low affinity values, respectively.

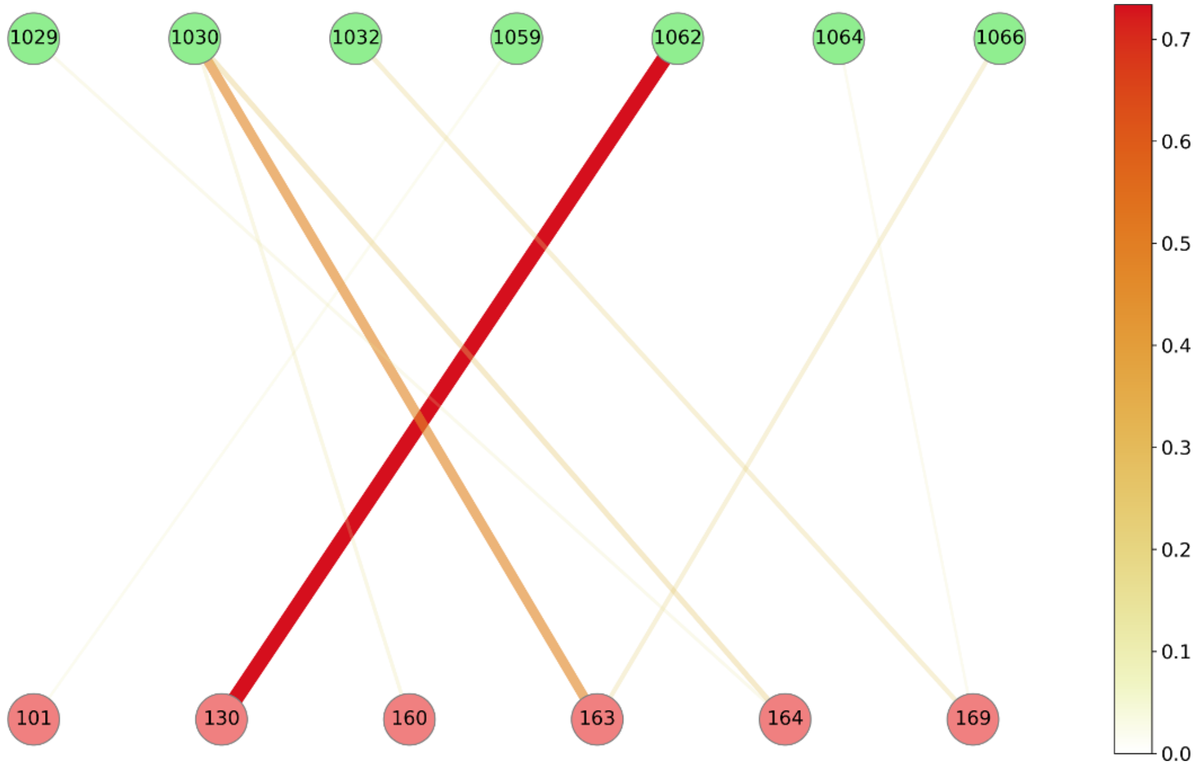


**Figure S13: Residue interaction network in final model of MP domain (red) with TSP7 (green) for cluster0 frames from Sim5.** Contacts of affinities below 0.15 were cut off. The connection line thickness and transparency are quadratically weighted by the affinities and colored from red to white for high and low affinity values, respectively.

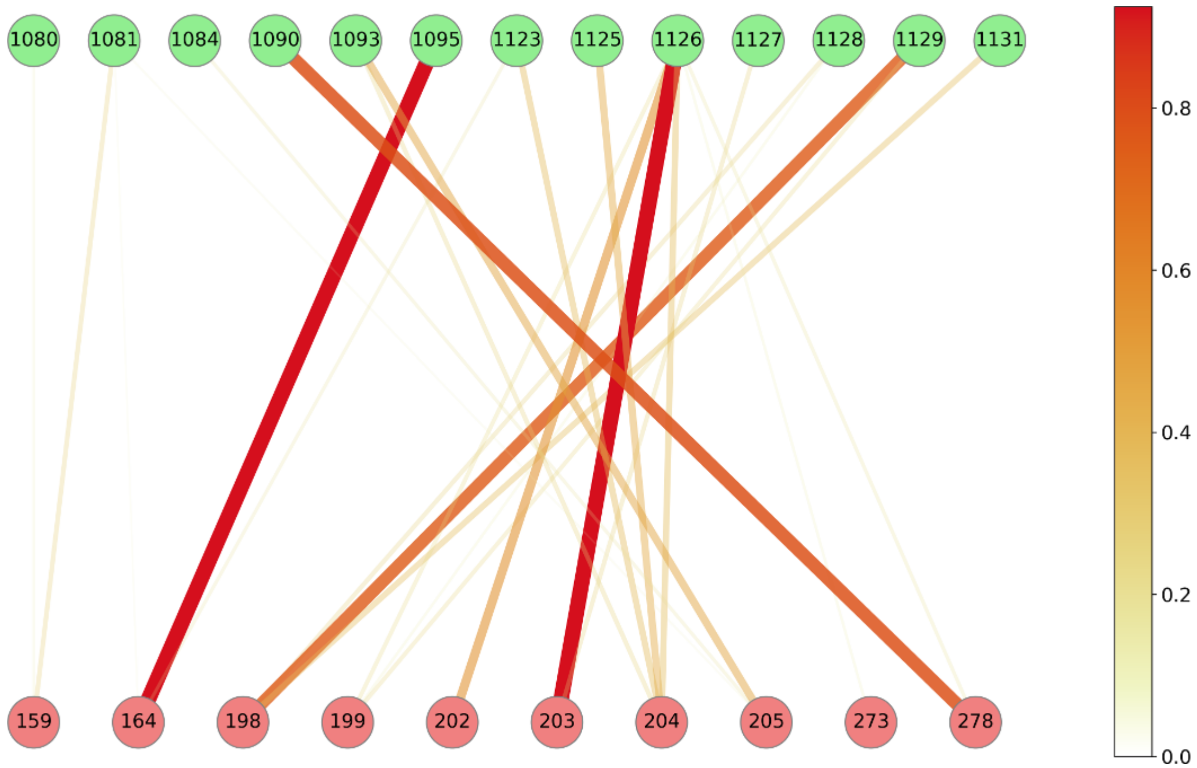


**Figure S14: Residue interaction network in final model of MP domain (red) with TSP8 (green) for cluster0 frames from Sim5.** Contacts of affinities below 0.15 were cut off. The connection line thickness and transparency are quadratically weighted by the affinities and colored from red to white for high and low affinity values, respectively.

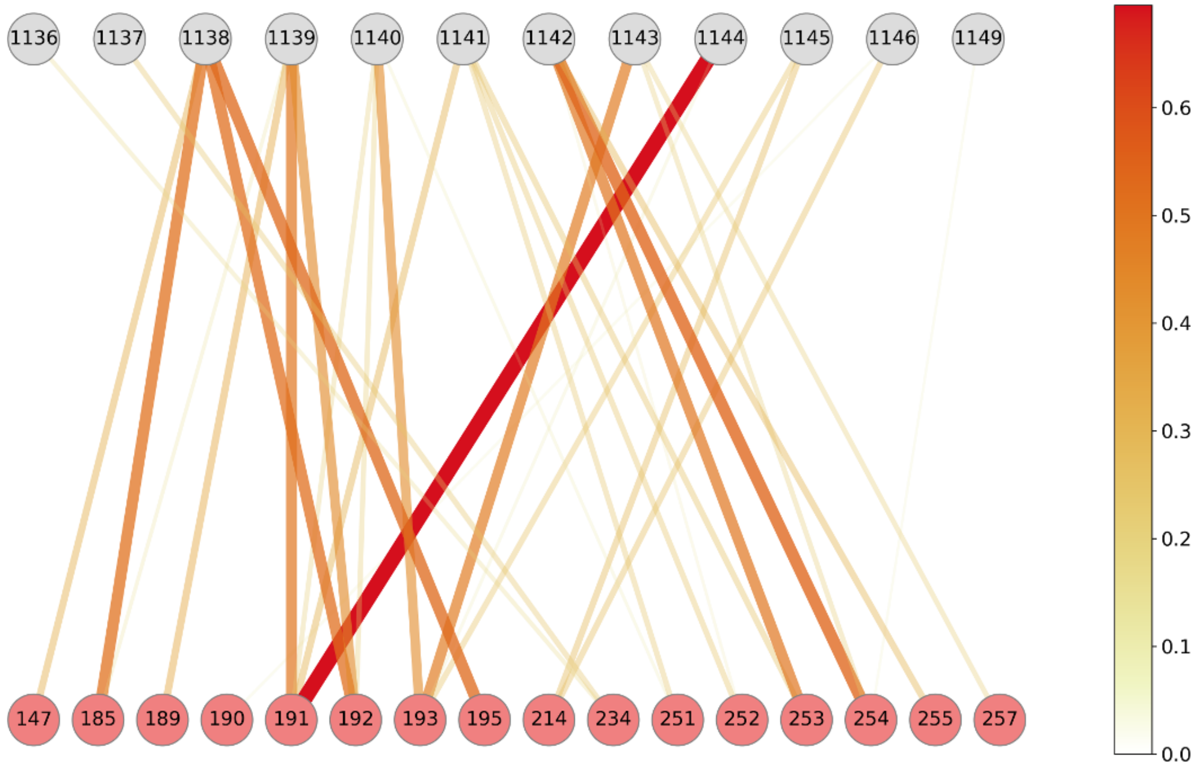


**Figure S15: Residue interaction network in final model of MP domain (red) with Linker (gray) for cluster0 frames from Sim5.** Contacts of affinities below 0.1 were cut off. The connection line thickness and transparency are quadratically weighted by the affinities and colored from red to white for high and low affinity values, respectively.

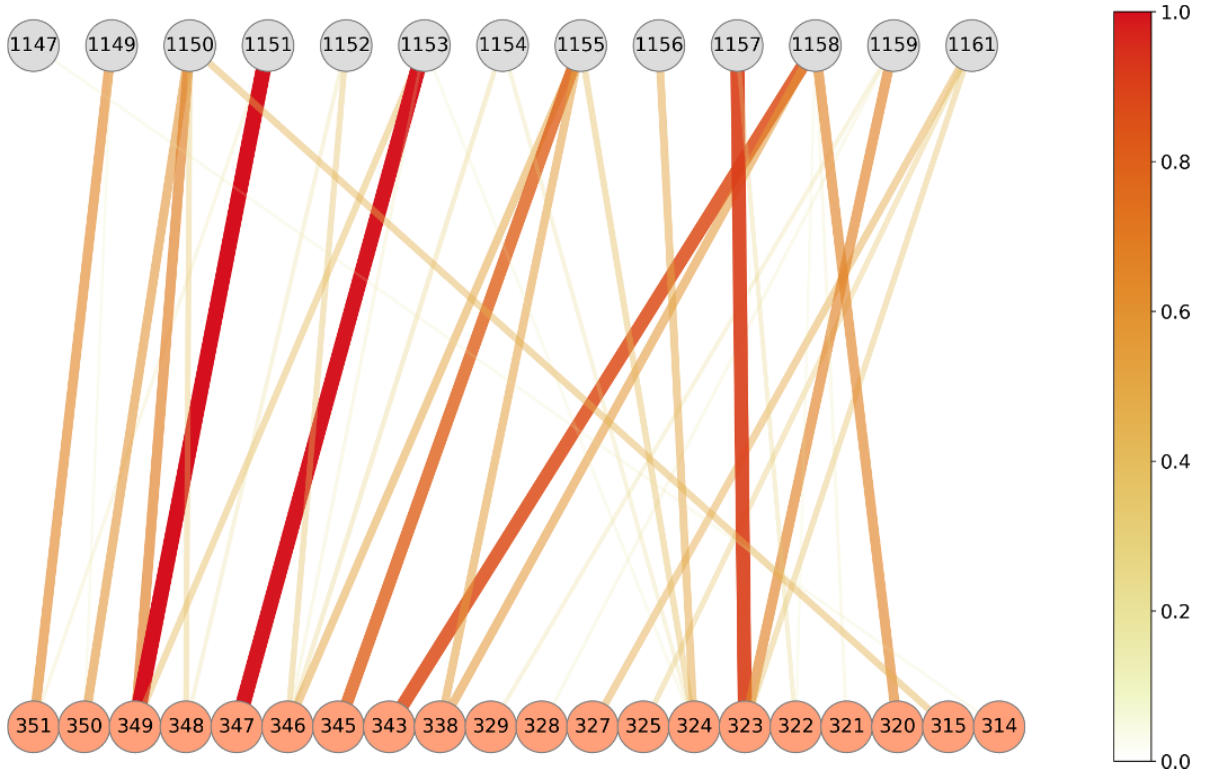


**Figure S16: Residue interaction network in final model of Dis domain (orange) with Linker (gray) for cluster0 frames from Sim5.** Contacts of affinities below 0.1 were cut off. The connection line thickness and transparency are quadratically weighted by the affinities and colored from red to white for high and low affinity values, respectively.

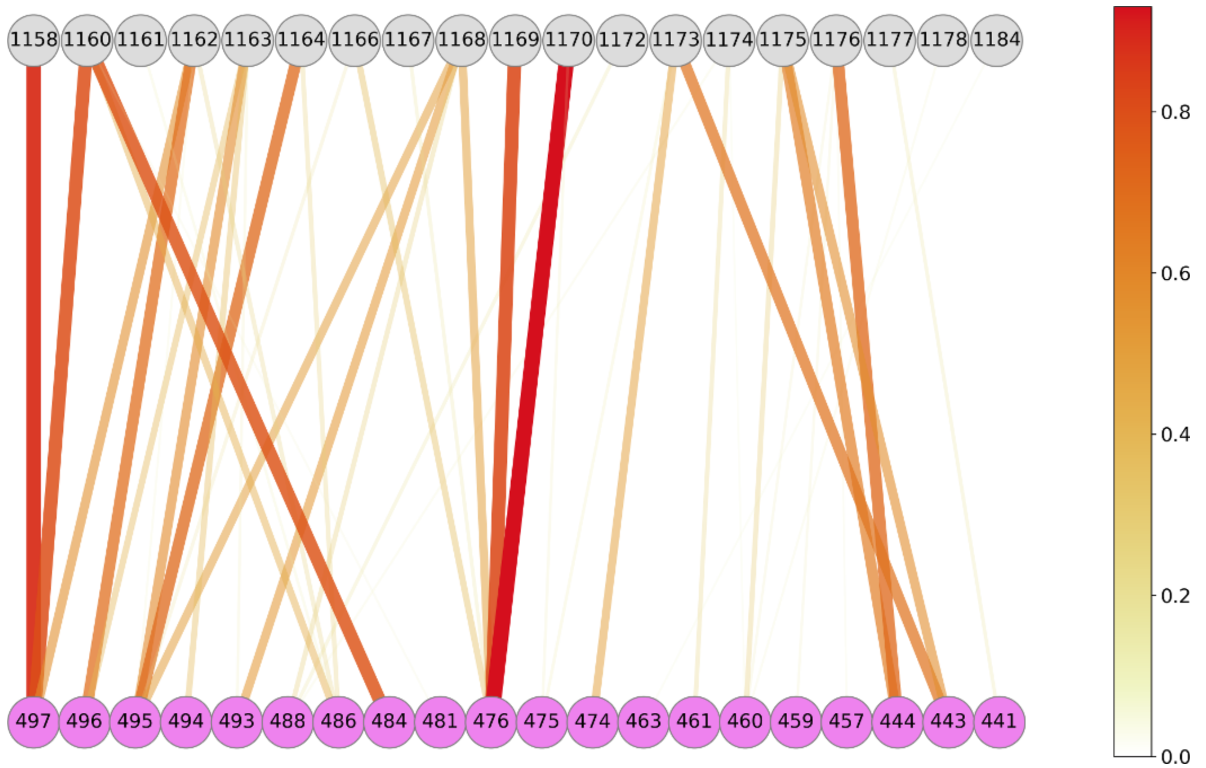


**Figure S17: Residue interaction network in final model of Cys domain (purple) with Linker (gray) for cluster0 frames from Sim5.** Contacts of affinities below 0.1 were cut off. The connection line thickness and transparency are quadratically weighted by the affinities and colored from red to white for high and low affinity values, respectively.

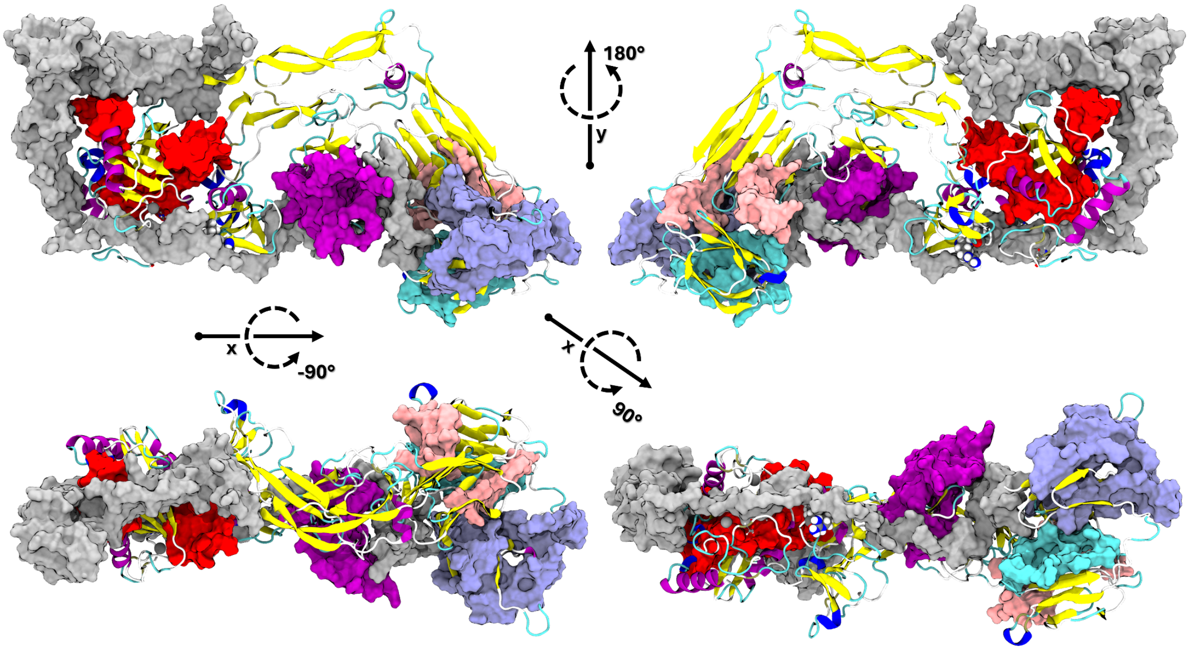


**Figure S18: HDX-MS results in agreement with final conformationally inactive ADAMTS13-Del3To6 model from Sim5 (cluster 0).** Gray surfaces represent TSP7, TSP8, and the Linker region, which obstruct the colored surfaces indicating HDX-MS results from Pillay *et al.*[24]. Highlighted residues correspond to regions in MP (red: 88-108, 158-174, 217-230), Cys-rich (purple: 446-482, 495-501), Spacer (pink: 629-642, 655-664), CUB1 (cyan: 1185-1214) and CUB2 (violet: 1313-1330, 1341-1347, 1358-1378, 1393-1407) domains that showed altered deuterium uptake in full-length compared with truncated ADAMTS13.
